# Dynamic proteomic and phosphoproteomic atlas of corticostriatal axon neurodevelopment

**DOI:** 10.1101/2022.03.21.485234

**Authors:** V. Dumrongprechachan, R. B. Salisbury, L. Butler, M. L. MacDonald, Y. Kozorovitskiy

## Abstract

Mammalian axonal development begins in embryonic stages and continues postnatally. After birth, axonal proteomic landscape changes rapidly, coordinated by transcription, protein turnover, and post-translational modifications. Comprehensive profiling of axonal proteomes across neurodevelopment is limited, with most studies lacking cell-type and neural circuit specificity, resulting in substantial information loss. We create a Cre-dependent APEX2 reporter mouse line and map cell-type specific proteome of corticostriatal projections across postnatal development. We synthesize analysis frameworks to define temporal patterns of axonal proteome and phosphoproteome, identifying co-regulated proteins and phosphorylations associated with genetic risk for human brain disorders. We discover proline-directed kinases as major developmental regulators. APEX2 transgenic reporter proximity labeling offers flexible strategies for subcellular proteomics with cell type specificity in early neurodevelopment, a critical period for neuropsychiatric disease.

## INTRODUCTION

Neurons are morphologically diverse cells that compartmentalize cellular signaling and information processing in cell bodies, dendrites, and axons, enabling spatiotemporal control over synaptic transmission. While these subcellular domains are maintained and regulated by both local and distal cues, local proteome regulation is particularly important for fine-tuning neural connectivity during circuit development and in the context of plasticity (Gonzalez-Lozano et al., 2016; Poulopoulos et al., 2019; Schanzenbächer et al., 2018). During the embryonic and early postnatal stages, axons rapidly grow and navigate to target regions across the brain (Winnubst et al., 2019). Developing axons require local protein synthesis and dynamics to coordinate axon outgrowth and growth cone collapse (Lin and Holt, 2007). Although axons contain machinery for local protein synthesis (Hafner et al., 2019), a large fraction of the neuroproteome is synthesized in the soma and trafficked throughout the cell, as evidenced by generally greater abundance of transcripts in the soma, compared to neurites (Glock et al., 2021). Thus, proteomic rather than transcriptomic techniques are necessary to capture the functional state of axons; this is especially important for axon guidance signaling that is highly regulated by post translational modifications such as protein phosphorylation (Costa-Mattioli et al., 2009). In addition, neuronal cell types are characterized by distinct patterns of gene expression (Saunders et al., 2018; Zeisel et al., 2018), implying biodiversity of their somatic, dendritic, and axonal proteomes. Therefore, it is important that measurements of the microdomain proteomes also contain cell-type specific information.

The postnatal proteome of developing axons remains largely unknown for any neuronal type. Mapping axonal proteomes from specific neurons directly in the vertebrate brain remains extremely challenging. Axonal compartments, such as axon growth cones, are typically isolated by tissue dissociation and sorting techniques, resulting in substantial losses of axon components (Chauhan et al., 2020; Poulopoulos et al., 2019). Axons from different circuits and cell types are intermingled, and isolation techniques that lack genetic targeting likely mask neuronal class specific features. Several genetically encoded proteomics strategies have been developed to profile cell-type specific proteomes in the mouse brain, including metabolic labeling (Alvarez-Castelao et al., 2017; Krogager et al., 2018) and proximity labeling strategies (Dumrongprechachan et al., 2021; Hobson et al., 2022; Rayaprolu et al., 2021; Spence et al., 2019; Uezu et al., 2016). We and others have previously shown that APEX-based proximity labeling in the acute brain sections is highly efficient and robust, providing sufficient material with both cell-type and subcellular compartment specificity such that there is no need for tissue dissociation, biochemical fractionation, or subject pooling (Dumrongprechachan et al., 2021; Hobson et al., 2022). APEX2 was engineered to be highly efficient for rapid protein biotinylation within seconds to minutes in the presence of hydrogen peroxides (Lam et al., 2015), providing superior labeling speed over other proximity labeling methods (Hung et al., 2016; Lobingier et al., 2017). Our prior work also demonstrates that rapid labeling speed of virally transduced APEX is capable of taking proteomic snapshots and capturing changes in the proteome within 3-4 hr time windows (Dumrongprechachan et al., 2021), but it is incompatible with applications that require early postnatal time points. Although APEX-based proximity labeling is a promising candidate for accessing the proteomic landscape of neurodevelopment, significant modifications to the overall strategy are needed in order to successfully map genetically targeted axonal proteomes in the mouse brain during postnatal development.

In this study, we focus on the postnatal development of corticostriatal projections. Corticostriatal systems represent a powerful model for studying axon development and axon guidance. In rodents, corticostriatal projections arise in the embryonic period and continue to innervate the striatum postnatally (Nisenbaum et al., 1998; Sheth et al., 1998; Sohur et al., 2014), where growth cones start to disappear around postnatal day P7, based on the reduction of growth-associated protein 43 (GAP43) immunostaining (Dani et al., 1991). As axon outgrowth declines after P7, the maturation of striatal neuron physiology becomes more pronounced, with high levels of dendritic spinogenesis, synaptogenesis, and tuning of intrinsic excitability until reaching maturity (P8 – P30) (Kozorovitskiy et al., 2012, 2015; Krajeski et al., 2019; Lieberman et al., 2018; Peixoto et al., 2016). Therefore, the dynamic phases of axonal development in the striatum raise a question about how the local axonal proteome changes across the first three postnatal weeks. Further understanding of typical neurodevelopment provides insight into how these processes break down in disease, as corticostriatal circuits are implicated in a number of neurodevelopment disorders such as autism spectrum disorder and schizophrenia. Because the striatum receives excitatory inputs from several brain regions (Hunnicutt et al., 2016), conventional approaches without a genetic targeting strategy cannot distinguish axons from different input types. To date, no genetically targeted proteome of axons across development has been reported. Furthermore, no prior study has succeeded in measuring genetically targeted phosphoproteome from any cell type in vertebrate brain. Phosphorylation state information, along with proteomics measurements, would provide an unparalleled window into local protein dynamics critical for neurodevelopment. Current APEX-based proteomics approaches primarily use Cre-dependent adeno-associated viral transduction (AAV) to deliver APEX reporters. This workflow restricts possible experimental timelines, because optimal expression of Cre-dependent AAV may take 1–3 weeks, rendering the early postnatal proteome intractable.

To address both technical limitations and biological questions in this study, we created a Cre-dependent APEX2 reporter mouse line using CRISPR knock-in. We demonstrated that the APEX reporter line can be used with multiple Cre-driver lines and is suitable for proteomics applications, including in the early postnatal period. We mapped the temporal expression patterns of the corticostriatal projection proteome during postnatal development. Combining APEX-based proximity labeling with phosphopeptide enrichment enabled a characterization of the local phosphoproteome in axons, revealing proline-directed kinases and phosphosites as major regulators for corticostriatal projection development.

## RESULTS

### Generation of a Cre-dependent APEX2 reporter mouse line

We created a Cre-dependent APEX reporter mouse line for broad applications in neuroscience and biology. A monomeric APEX2 variant was chosen because of its small size and efficient activity for proximity labeling of proteins (Lam et al., 2015). A double-floxed inverted orientation of the APEX transgene (DIO-APEX2-P2A-EGFP) was introduced into the ROSA26 locus under the EF1a promoter using CRISPR/Cas9 strategy (**Figure 1A**). Zygotes were microinjected with enhanced specificity eSpCas9, HDR template and crRNA:tracrRNA duplex, using a previously validated gRNA sequence targeting the ROSA26 locus (Chu et al., 2016). We identified 8 out 112 founder mutants by genotyping PCR (**Figure 1B, Table S1**). We confirmed germline transmission and mendelian inheritance by crossing selected founders to C57BL/6 wildtype mice over two generations. The knock-in line was bred to homozygosity by crossing F2 heterozygotes (**Figure 1C**). We evaluated transgene integration using long-range PCR amplification. Using primers inside the transgene and outside the left or right homology arms, respectively, we found that the size of PCR amplicons matched the predicted sequences, suggesting that transgene integration was site-specific (**Figure 1D**). In addition, we PCR amplified and sequenced top eight predicted off-target sites with NGG PAM across 6 founders. We found a small number of off-targets that were affected by Cas9 activity on non-coding protein regions (**Figure S1A–C, Table S2**). We did not observe noticeable phenotypes in either heterozygous or homozygous animals. From this point onward, we chose to work with founder line #83 (**Figure S1A–C**). To evaluate conditional expression of the APEX transgene, the APEX reporter line was crossed to VGlut2-Cre or VGAT-Cre mice (Vong et al., 2011). We performed direction-specific PCR to assess Cre-dependent inversion of the APEX transgene using genomic DNA that was directly extracted from the cortex. Inverted transgene was only detected in Cre-expressing brain tissues (**Fig. 1E**), confirming Cre-dependent recombination. We also inspected the pattern of EGFP reporter expression across multiple brain regions in both VGlut2-Cre and VGAT-Cre crosses (**Fig. 1F–G**). Dense reporter expression was observed in the cortex for VGlut2-Cre and in the striatum for VGAT-Cre, respectively. In the hippocampus, the granule cell layer in the dentate gyrus was clearly visible in VGlut2-Cre cross, with smaller numbers of GABAergic cells present in the hilus of VGAT-Cre cross. In the cerebellum, Purkinje cell layer was observed in VGAT-Cre and absent in VGlut2-Cre mice crossed to the APEX reporter. Both genetic and histological evidence demonstrates Cre-dependent control of the APEX transgene.

**Figure 1.**
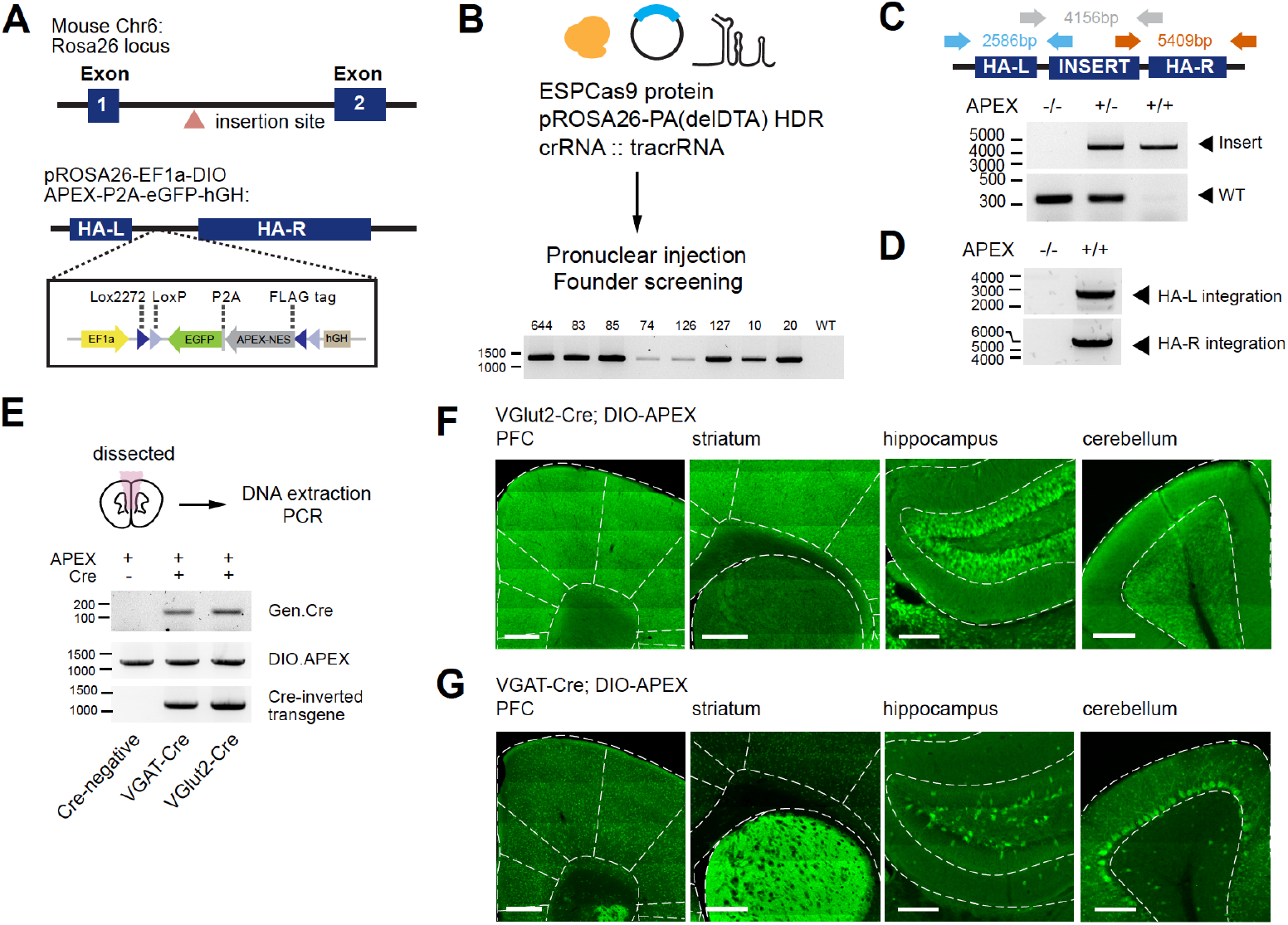
Generation of a Cre-dependent APEX2 reporter mouse line. **(A)** Design of a Cre-dependent APEX reporter mouse line. A Cre-dependent APEX transgene under the control of EF1a promoter is targeted for knock-in at the Rosa26 locus in the mouse genome. Inverted APEX-NES (nuclear exporting sequence) is flanked by lox sites in a double floxed inverted orientation (DIO). **(B)** CRISPR/Cas9 strategy for mouse line generation. Enhanced specificity Cas9 (ESPCas9), targeting vector, and crRNA::tracrRNA duplex were used in pronuclear injection to generate founders. Agarose gel electrophoresis shows founders containing APEX transgene. **(C)** PCR amplification across the APEX transgene. Gray arrows: 317 bp WT loci and ~4 kb APEX transgene. **(D)** PCR amplification across the homology arms evaluating genomic integration. Blue arrow, left homology arm (HA-L), 2586 bp. Red arrow, right homology arm (HA-R), 5409 bp. **(E)** PCR amplification of genomic DNA extracted from the brain for the presence of Cre, the APEX transgene before and after Cre-mediated recombination in Cre-/-, VGAT-Cre+/- and VGlut2-Cre+/- animals. **(F)** Cre-dependent expression of EGFP reporter in the prefrontal cortex (PFC), striatum (STR), hippocampus (HC), and cerebellum (CB) of VGlut2-Cre mice crossed to APEX reporter line (immunostaining using anti-GFP antibody). Scale bars, 500 μm for PFC and STR, and 200 μm for HC and CB. **(G)** same as **(F)** but for VGAT-Cre. See also **Figure S1**.

### Ex vivo biotinylation workflow for cell-type specific proteomics

To broadly distribute APEX inside neurons, we included the nuclear exporter sequence (NES) at the C-terminus of the APEX protein. We confirmed APEX localization using anti-FLAG immunofluorescent staining (**Figure 2A**). APEX localizes in the cytosolic compartment compared to untargeted EGFP structural markers, separated by a P2A linker. To demonstrate the utility of the APEX reporter line for proteomics, we used biotin phenol (BP) and H_2_O_2_ to induce biotinylation of proteins in APEX expressing neurons in the VGlut2 and VGAT crosses. We modified published APEX-mediated biotinylation protocols to optimize protein labeling in thick brain tissue (Dumrongprechachan et al., 2021), since APEX has been largely characterized and applied in cell culture (Hung et al., 2016; Lobingier et al., 2017; Paek et al., 2017) and small organisms (Chen et al., 2015; Reinke et al.). Our prior work and other studies have demonstrated that AAV-mediated APEX expression and biotinylation in the mouse brain is sufficient for cell-type specific mass spectrometry-based (MS) proteomics (Dumrongprechachan et al., 2021; Hobson et al., 2022). To label proteins in APEX-expressing tissues, acute brain slices of 250 μm thickness were prepared from VGlut2-Cre;APEX animals and incubated in ACSF supplemented with 500 μM BP up to 1hr. Following BP incubation, slices were briefly washed in ACSF and biotinylation was induced by 0.03% H_2_O_2_ treatment for 2 min and quenched with Na-ascorbate ACSF (**Figure 2B**). Streptavidin dot blot analysis of cortical protein lysates shows increasing amounts of biotinylated proteins over 60 min (**Figure 2A**). Western blot analysis of total prefrontal cortex (PFC) lysates at 60 min BP incubation reveals differential patterns of protein labeling in VGlut2-Cre and VGAT-Cre, with negligible labeling in the Cre-negative APEX controls (**Figure S2B**). As expected, the level of biotinylation in VGlut2 PFC sample was greater than that of VGAT, consistent with a greater number of glutamatergic neurons relatively to GABAergic neurons present in the neocortex, since under ~20% of cortical neurons are GABAergic (Keller et al., 2018). Using the 1 hr-long incubation protocol, we confirmed that biotinylation is both Cre and APEX-dependent (**Figure 2C**). We observed some labeling in VGlut2-Cre+ samples during 1-hr BP incubation without H_2_O_2_, and labeling efficiency was greatly enhanced in the presence of H_2_O_2_. To streamline sample preparation for MS-based proteomics, we evaluated the enrichment method using streptavidin magnetic beads (**Figure 2D**). Successful enrichment was verified by the depletion of streptavidin signal in flowthrough fractions and the presence of high streptavidin signal in the enrichment output, where the corresponding total amount of enriched proteins is shown by silver staining. Altogether, these experiments establish a method for selective proteome labeling, which we next apply to profile the proteome of excitatory cortical inputs to the striatum across postnatal development.

**Figure 2.**
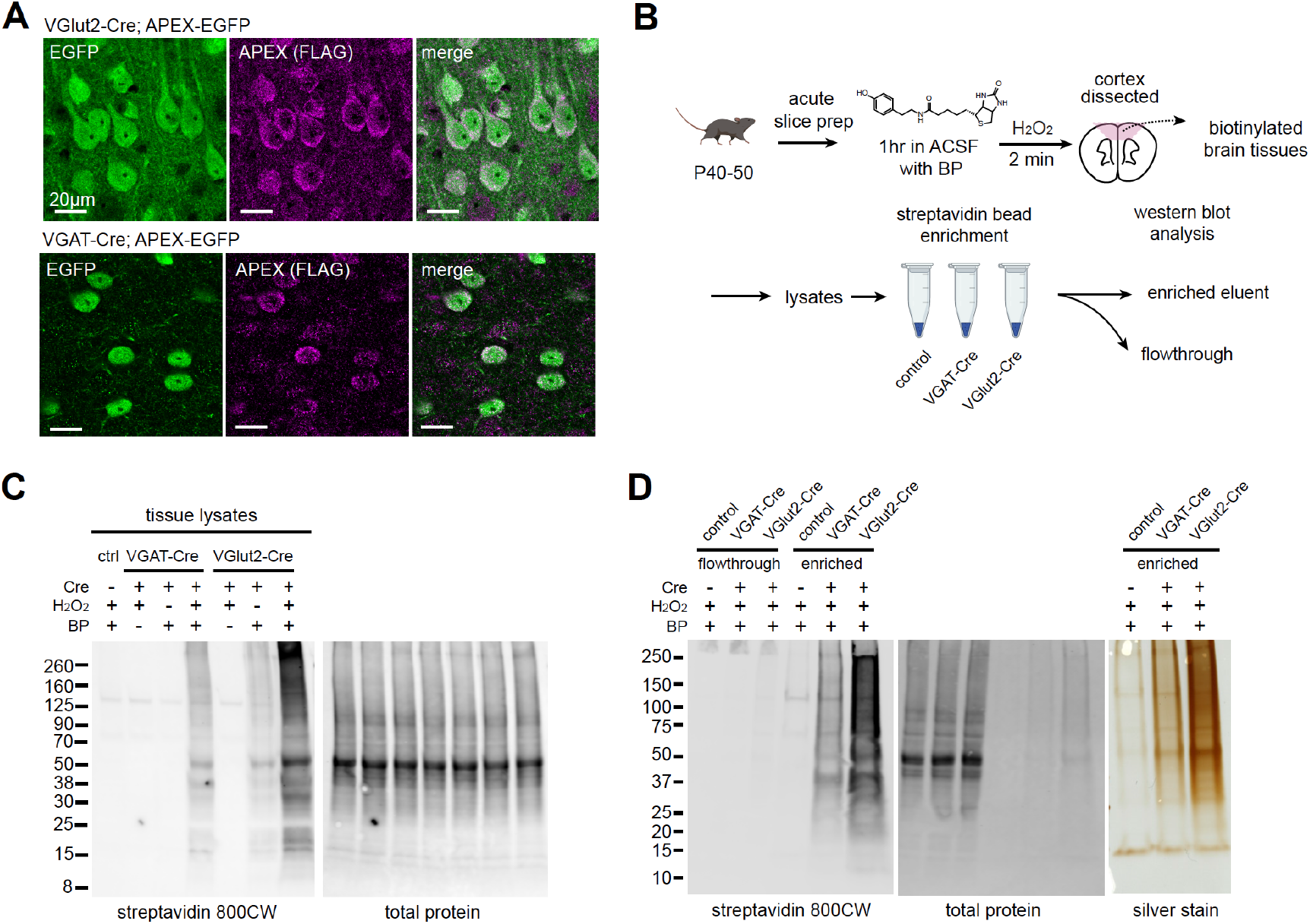
APEX-mediated biotinylation in brain tissue. **(A)** APEX subcellular localization in the mPFC of VGlut2-Cre;APEX and VGAT-Cre;APEX (immunostained EGFP 488, anti-FLAG 647 for APEX, scale bars 20 μm). **(B)** Sample preparation schematic for protein analysis. Acute slices were incubated in carbogenated artificial cerebrospinal fluid (ACSF) supplemented with 500 μM biotin phenol for 1 hr. Sections were briefly rinsed in ACSF. APEX was activated by 0.03% H_2_O_2_ ACSF for 2 min and quenched in Na ascorbate ACSF solution. Protein lysates were prepared from dissected tissues. Streptavidin beads were used to enrich tagged proteins for analysis. **(C)** Cell-type specific biotinylation in Cre-negative, VGAT-Cre, and VGlut2-Cre crosses. Western blot analysis of cortical protein lysates. Biotinylated proteins and total protein loading control were detected by streptavidin-CW 800 and REVERT 700 stain, respectively. **(D)** Validation of streptavidin bead enrichment. *Left*, detection of biotinylated proteins in flow-through and enriched fractions. *Right*, silver stain gel of enrichment output proteins. Differential enrichment outputs reflect differences in GABAergic and glutamatergic cell proportions in the cortex. See also **Figure S2**.

### Selective biotinylation of corticostriatal axons across postnatal development

Glutamatergic inputs into the striatum develop rapidly across embryonic and early postnatal stages (Kuo and Liu, 2019; Peixoto et al., 2016), yet deep knowledge of the overall proteome dynamics in developing cortico-striatal projections has remained out of reach for technical reasons. To capture axonal proteome dynamics during this process, we crossed the Rbp4^Cre^ mouse line to our APEX reporter (Gerfen et al., 2013). As expected, Rbp4^Cre^;APEX expression is primarily restricted to layer 4–5 corticostriatal projection neurons (**Figure 3A**). We showed that EGFP reporter expression is observed in the striatum throughout postnatal time period P5-P40 (**Figure 3A**). Western blot analysis after ex vivo biotinylation detects labeled proteins in both cortical and striatal tissues above the negative control, including at the earliest P5 time point (**Figure 3B, Figure S3A**). This is especially important because viral based genetically encoded proteomics is limited for early development applications (**Figure S3B**). No somatic expression was observed in the striatum; therefore, biotinylated proteins in the striatal lysates represent corticostriatal axonal proteins. Having validated axonal targeting strategy, we used mass spectrometry-based bottom-up proteomics to identify and quantify the axonal proteome of the striatum. In addition, to rigorously evaluate the extent of axonal enrichment for each protein, we included a somatic compartment control in the experiment. We used adeno-associated viral transduction to express nuclear localized H2B.APEX broadly in the cortex of Rbp4^Cre^ mice. Immunofluorescence (**Figure S3C–D**) and western blot analysis (**Figure S3E–F**) show that protein biotinylation in H2B.APEX-expressing Rpb4Cre animals is restricted to the cortex and absent in the striatum. Thus, H2B.APEX is an optimal somatic compartment control because its labeling is primarily restricted to the nucleus and soma.

**Figure 3.**
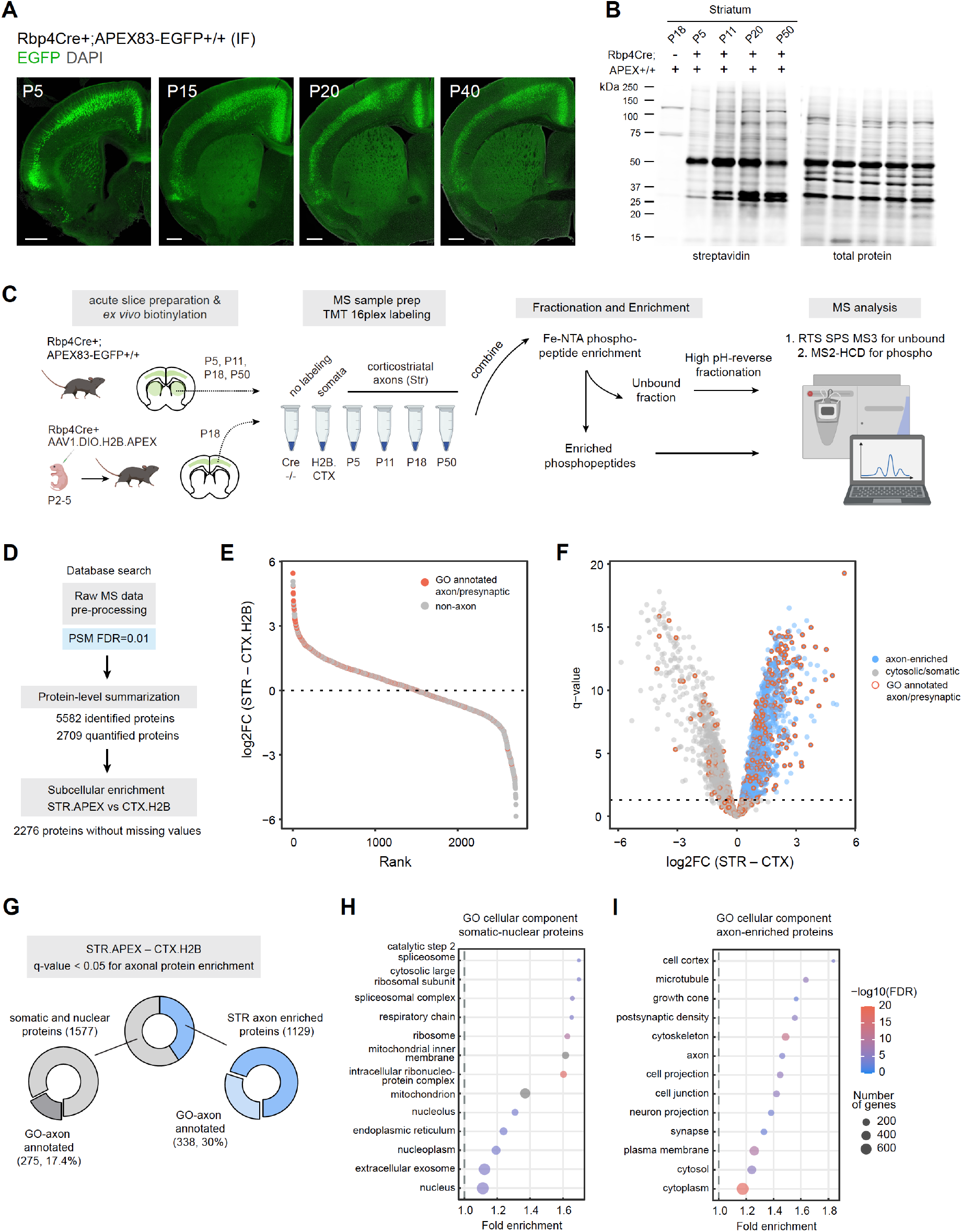
Genetic targeting approach for proteomics analysis of corticostriatal axonal compartment. **(A)** Cre-dependent APEX.EGFP expression in excitatory cortical inputs to the striatum. Expression pattern of APEX.EGFP (immunostained EGFP) in Rbp4^Cre+^;APEX-EGFP+/+ animals across postnatal days 5, 15, 20, and 40. Coronal brain section, scale bar: 1 mm. **(B)** Biotinylation of proteins in corticostriatal axons across development. Western blot analysis of striatal lysates prepared from biotinylated acute brain slices. Acute slices were prepared and biotinylated from Rpb4^Cre+^;APEX animals at the indicated age. Cre-negative APEX reporter line at P18 was used as no labeling control. *Left*, streptavidin blot. *Right*, total protein loading control. Lower amount of protein biotinylation at early age (P5) is consistent with developing innervation by axonal projections (see **Figure 3A**). **(C)** Sample preparation workflow for proteomics analysis. Acute slices were prepared and biotinylated from Rbp4^Cre+^;APEX or Rbp4^Cre+^ injected with H2B.APEX at the indicated time points. Streptavidin beads were used to enrich biotinylated proteins for the bottom-up proteomics workflow. Samples were labeled with tandem mass tags (TMT) and combined. TMT mixtures were enriched for phosphopeptide analysis using Fe-NTA tips. The unbound fractions were fractionated using high pH-reverse phase resin for protein abundance analysis. Synchronous precursor selection (SPS) MS3 method was used for proteome analysis. MS2 with high energy collisional dissociation (HCD) method was used for phosphopeptide analysis. **(D)** Data analysis flow chart. Raw MS data were analyzed by Proteome Discoverer with false discovery rate (FDR) set at 1% for peptide spectrum matches (PSMs). Protein-level summarization was performed using MSstatsTMT using unique peptides. Striatal (STR) and cortical (CTX.H2B) samples prepared from P18 were compared to determine the axon-enriched proteome. Proteins with missing values were removed prior to this comparison. **(E)** Log2-fold change (FC) plot for STR–CTX comparison. Proteins were ranked by their log2FC. Red and gray dots, proteins with or without presynaptic or axon gene ontology annotations, respectively. **(F)** Volcano plot for the STR–CTX comparison. Blue dots, axon-enriched proteins with log2FC > 0 and q-value < 0.05 (horizontal dotted line). Gray dots, somatic or nuclear proteins that did not pass axon enrichment cutoff. Red border, proteins with presynaptic or axon gene ontology annotations. **(G)** Proportion of proteins with GO axon/presynaptic annotation for somatic/nuclear and axon protein lists. Using STR–CTX comparison, the proteome is classified into somatic/nuclear and axon-enriched proteins. **(H)** Over-representation analysis using gene ontology (GO) cellular component database for somatic and nuclear enriched protein list. Dot plot ranked by fold enrichment. Dot size and color correspond to the number of genes, and −log10(FDR), respectively. **(I)** Same as **(H)**, but for the axon-enriched protein list. See also **Figure S3**.

In our experimental design (**Figure 3C**), we biotinylated and enriched axonal proteins from the striatum across 4 time points (~P5, P11, P18, and P50, n = 5 biological replicates) (**Table S3**). We targeted these early time points, because striatal circuits start to transition from axon innervation to synaptogenesis, which lead to maturation of striatal neurons as animals enter adolescence and adulthood (Kuo and Liu, 2019). Additionally, we included samples prepared from the Rbp4^Cre+^ cortex expressing H2B.APEX as the somatic compartment control (n = 3) from P18 animals. No labeling controls were collected from Cre-negative cortical and striatal samples that underwent the same ex vivo biotinylation procedure (n = 3, 4 for cortex and striatum replicates, respectively). Enriched biotinylated proteins were digested on-bead, and peptides were barcoded with tandem mass tag (TMT) for mass spectrometry analysis. TMT-labeled peptides were mixed equally and enriched for phosphopeptides. The unbound flowthrough was fractionated using the high-pH reverse fractionation to measure protein abundance. We adapted our previous analysis workflow using MSstatsTMT for protein summarization, and comparison with Cre-negative samples for removing non-specific binding contaminants (Dumrongprechachan et al., 2021). MSstatsTMT was used to summarize peptide intensity into protein abundance (Huang et al., 2020). We identified 5582 proteins and quantified 2276 out of 2709 proteins without missing values across conditions (**Figure 3D**) (**Data S1**). Boxplot of normalized protein abundance shows aligned medians across samples (**Figure S4A**). Principle component analysis (PCA) shows a distinct separation of sample groups with minimal TMT plex effects (**Figure S4B**).

To evaluate the extent of axonal enrichment for each protein, we used moderated t-tests implemented in MSstatsTMT to compare P18 striatal samples to the P18 H2B.APEX nuclear/somatic controls. We rank proteins by their log2 fold change (STR–CTX.H2B) (**Figure 3E**) (**Data S2**). Proteins with greater log2FC are more likely to have prior axon/presynaptic gene ontology annotations, consistent with a greater GO-axon term annotation rate as rank increases (**Figure S4C**). Proteins with q-value < 0.05 and log2FC > 0 were considered axonally enriched, depicted in the volcano plot (**Figure 3F**). Using this criterion, we loosely classified axonal proteins in the dataset into 1129 axon-enriched proteins and 1577 somato-nuclear proteins, with 30% and 17.4% having GO-axon annotations, respectively (**Figure 3G**). Overrepresentation analysis using the cellular component GO database shows an enrichment of cell projection-related terms for the axon-enriched proteome (*e.g*., axon, neuron projection, synapse) and nucleus and mitochondrion-related terms for the somato-nuclear proteome (*e.g*., cytosolic ribosomal subunits, mitochondria, nucleoplasm) (**Figure 3H–I**) (**Data S3**). Together, rank based analysis confirms that our approach successfully enriched axonal proteome and GO analysis breaks down the overall subcellular compartments present in the axons. This demonstrates a workflow for mapping axonal proteomes using the Cre-dependent APEX reporter mouse line.

### Temporal trajectories of the corticostriatal axonal proteome

Next, we evaluate how the axonal proteome changes across postnatal development. We stringently include only the proteins that were quantified in all biological replicates in this analysis. First, we globally assess whether protein is significantly altered across time, using the least-squares linear regression implemented in the maSigPro R package (Conesa et al., 2006). Neonate (P5) was chosen as the reference time point. maSigPro reported 2274 proteins with corresponding adjusted p-values (**Data S4**). As expected, the majority of the detected proteome was highly dynamic, with 1981 out of 2274 proteins significantly changing (adjusted p-values < 0.05). To better observe protein expression patterns, we performed hierarchical clustering of protein abundance, based on time points, revealing 8 general developmental trajectories (**Data S4**). The heatmap (**Figure 4A**) shows individual protein abundance normalized to the neonatal time point for each cluster, reflecting the dataset quality with overall CV < 10% (**Figure S5A**). For each cluster, we compute and plot the mean protein abundance (**Figure 4B**). Clusters 1, 2, 4, and 5 show increase protein expression over time, while clusters 3 and 8 are characterized by decreased protein expression over time. Clusters 6 and 7 fluctuate around the baseline with negative and positive peaks during the early postnatal time point, respectively. For quality control, we plotted several examples of known developmentally regulated proteins (Gonzalez-Lozano et al., 2016), including doublecortin (*Dcx*), neurofilament medium polypeptide (*Nefm*),synaptophysin (*Syp*), VGlut1 (*Slc17a7*), and vesicle marker *Rab3a* (**Figure S5B**). As expected, an immature neuronal marker like microtubule-associated protein *Dcx* rapidly decreases during development, while mature neuronal markers such as medium intermediate filament (*Nefm*) and other synaptic and vesicle markers (*Syp*, *Slc17a7*, and *Rab3a*) increase over time.

**Figure 4.**
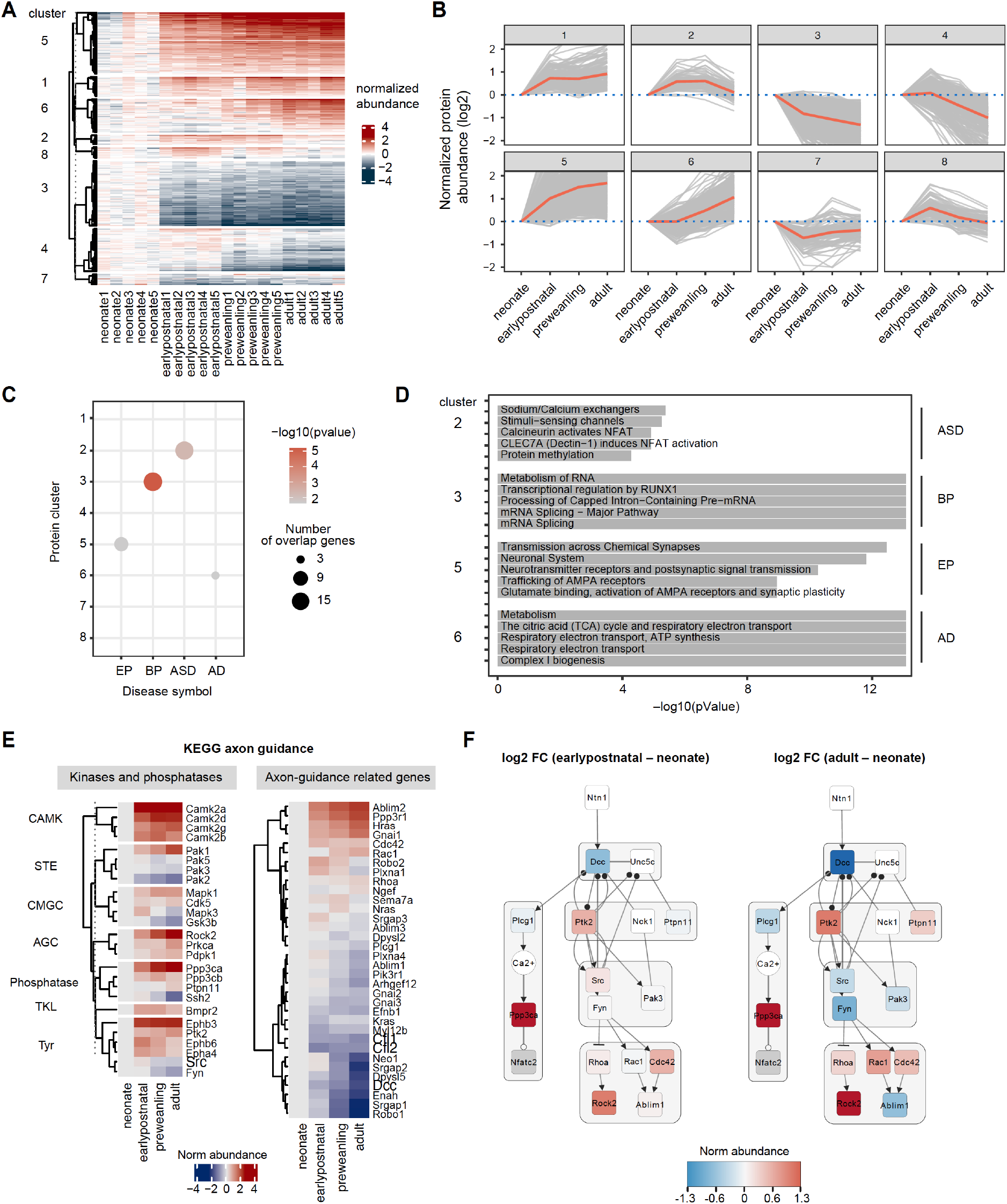
Temporal trajectories of the corticostriatal axonal proteome. **(A)** Heat map showing temporal expression across development. Protein clusters were defined using a time course regression analysis implemented in maSigPro R package. The row dendrogram shows hierarchical clustering of proteins in each protein cluster. Log2 protein abundance was normalized to the mean of the neonatal time point. All biological replicates are shown in columns. **(B)** Temporal trajectories of protein expression in each cluster. Log2 protein abundance was normalized to the mean of the neonatal time point. Red lines represent the average expression of all proteins in a cluster. Gray lines represent individual protein trajectories. Neonate (P5), early postnatal (P11), preweanling (P18–20), and adult (P50). **(C)** Over-representation analysis of protein clusters for central nervous system (CNS) traits and disorders. Dot size indicates the number of genes enriched for a particular disease. Color density indicates the degree of significance, −log(pvalue). Only significantly enriched diseases are shown. AD (Alzheimer’s disease), EP (Epilepsy), BP (bipolar disorders), and ASD (Autism spectrum disorders). **(D)** Reactome pathway enrichment analysis for protein clusters that are associated with CNS traits and disorders (clusters 2, 4, 7, and 8). Top 5 Reactome pathway terms are plotted. Gray bar indicates −log(p value). **(E)** KEGG axon guidance pathway analysis. Heatmap of normalized Log2 protein abundance for proteins mapped to the KEGG axon guidance pathway (mmu04360). *Left*, protein abundance for kinases and phosphatases clustered by family. *Right*, other proteins in the pathway. **(F)** Temporal changes in protein abundance in the Ntn1-Dcc signaling subnetwork between early postnatal and adult. Node color, Log2 fold change normalized to the neonate condition. Edges, solid and blunt arrows for activation and inhibition, closed and opened circles for phosphorylation and dephosphorylation, respectively. See also **Figure S4–5**.

Abnormal neurodevelopmental processes including axonal development have been implicated in many psychiatric diseases, with numerous genetic variations and mutations associated with specific disorders (Werling et al., 2020). While transcript levels in brain tissues from multiple species have been well cataloged across development (Colantuoni et al., 2011; Werling et al., 2020), the temporal patterns of protein and protein phosphorylation levels in neuronal subtypes are not fully mapped (Carlyle et al., 2017; Gonzalez-Lozano et al., 2016), which limits our understanding of how disease risk genes contribute to disease etiology. This question is especially important for our understanding of neuropsychiatric diseases of polygenic origin, where protein networks are believed to serve as the point of convergence for multiple genetic risk factors. To determine whether protein clusters are enriched for disease risk genes, we compiled a list of risk genes associated with 11 disorders: autism spectrum disorders (ASD), Alzheimer’s disease (AD), Parkinson’s disease (PD), epilepsy (Ep), developmental delay (DD), Schizophrenia (Scz), multiple sclerosis (MS), major depressive disorder (MDD), bipolar disorders (BP), attention deficit hyperactivity disorder (ADHD), and glioma (**Data S5**) (Abrahams et al., 2013; Chang et al., 2017; Consortium et al., 2020; Deciphering Developmental Disorders Study, 2017; Demontis et al., 2019; Fu et al., 2021; Heyne et al., 2018; International Multiple Sclerosis Genetics Consortium, 2019; Rice et al., 2016; Stahl et al., 2019; Wightman et al., 2021; Wray et al., 2018). We used hypergeometric testing implemented in Webgestal (Wang et al., 2017) to statistically evaluate risk gene enrichment using the 2107 maSigPro quantified proteins as the background. Clusters with p-value < 0.5 and FDR < 0.3 were considered significant. We found that clusters 2, 3, 5, and 6 were enriched in ASD, BP, EP, and AD risk genes, respectively (**Figure 4C**, **Data S6**). In addition, we performed Reactome pathway enrichment analysis (Jassal et al., 2020) to look at which cellular pathways associate with each protein cluster (**Figure 4D**, **Data S6**). We examine the relationship between risk genes and biological pathways in the cluster using STRING-DB interactome analysis (**Figure S5C**). For example, we found that epilepsy risk genes are significantly enriched in protein cluster 5 (p-value = 0.022), which is composed of proteins with increasing expression over time. This implies that postnatal increase in these gene products is important for proper neural circuit development. Cluster 5 also contains genes enriched in α-amino-3-hydroxy-5-methyl-4-isoxazolepropionic acid receptor (AMPAR) trafficking and plasticity (p = 1.1e-9), linking epilepsy risk genes in this cluster to excitatory glutamatergic transmission. In fact, reduced corticostriatal excitatory transmission to fast-spiking interneurons was found to trigger absence seizures in Stxbp1 haplodeficient mouse model and this effect was mitigated by pharmacological activation of AMPAR in striatum (Miyamoto et al., 2019). Altogether, this analysis framework generates an overview of how cellular pathways, developmental trajectories, and genetic risk for neuropsychiatric diseases relate to one another in the context of normal axon development in the striatum.

### Pathway-centric analysis for the axon guidance system

To gain deeper insights into the regulation of axon development, we performed pathway-centric analyses focusing on signaling pathways during axon guidance. We mapped our proximity labeling axon proteome data to the KEGG axon guidance pathway (Kanehisa et al., 2017). Because kinases and phosphatases are the major regulators of axonal development, we classified mapped genes into two categories: kinases/phosphatases, and axon-guidance related genes. Temporal trajectories of KEGG genes were visualized in heatmaps with row clustering, separating kinases and phosphatases by their families (**Figure 4E**). We found that the majority of kinases increase in expression over time, such as CAMK and AGC families. However, some kinases (*e.g., Pak2, Pak3, Pak5, Gsk3b, Ssh2, Src*, and *Fyn*) show elevated expression during neonatal and early postnatal time points, but decrease in expression over time, suggesting a functional relevance for early axon guidance. In addition, we detected kinases that are known for their crucial roles in neurodevelopment, including the *Mapk* family, *Gsk3b, Cdk5*, and *Src/Fyn. Mapk, Gsk3b*, and *Cdk5* are proline-directed kinases while *Src* and *Fyn* are non-receptor tyrosine kinases. As for other axon guidance related KEGG genes, we can group them by their functional similarities including ligand-guidance receptor systems, GTPases and their regulators, and cytoskeleton remodeling proteins. In particular, small GTPases including *Rhoa, Rac1*, and *Cdc42* are important actin cytoskeleton regulators that control axon growth cone expansion during development (Shekarabi et al., 2005).

As an example of mining the datasets generated here, we show the Netrin1 (*Ntn1*)-*Dcc* subnetwork in the KEGG axon guidance pathway to investigate whether corticostriatal projections follow a canonical model of axon development (**Figure 4F**). *Dcc* is a netrin receptor that expresses in the axon (Vosberg et al., 2020). Activation of the *Dcc* pathway by netrin promotes axon outgrowth by recruiting regulators of actin cytoskeleton (Shekarabi et al., 2005). This process depends on *Fyn*-mediated phosphorylation of *Dcc* (Meriane et al., 2004) as well as *Src*-mediated phosphorylation of *Plcg1* (Kang et al., 2018). Our data show that *Dcc* expression decreases over time, reflected by negative log2 fold change in the early postnatal and adult samples compared to the neonate. Similarly, *Src* and *Fyn* kinases are upregulated during the early postnatal period and downregulated in the adulthood. The timing of *Src* and *Fyn* upregulation in the early postnatal time coincides high *Dcc* expression, implicating *Src* and *Fyn* as primary regulators of *Dcc* signaling in the neonatal and early postnatal time windows, but not in the adult with matured striatal circuits. Our data for corticostriatal systems are in an agreement with previous reports examining axonal *Dcc* signaling in Xenopus laevis retinal ganglion cells (Meriane et al., 2004) and in mouse midbrain dopaminergic neurons (Kang et al., 2018), validating these data as an important resource for evaluating axonal proteomes.

### Phosphosite-centric analysis of kinase substrate interactions

Measuring protein levels in time and space offers many clues as to how a biological system matures across development and how it can break down in neuropsychiatric disease. However, it cannot measure protein state (*e.g*., activity) which is regulated in large part by post translational modifications. Even the abundance of protein kinases alone is insufficient to infer their activity and their constellation of downstream phosphorylation; thus, a direct measurement of protein phosphorylation is essential. This is especially important in the context of axon development, where coordinated phosphorylation events control axon pathfinding, growth cone elongation, and growth cone collapse. Therefore, during the study design, we aimed to create a workflow that enables quantification of both protein and phosphopeptide abundance from the same samples (**Figure 3C**). During sample preparation, on-bead digested peptides were labeled with TMTPro reagents, desalted, and then phospho-enriched. We found that TMTPro multiplexing provides sufficient phosphopeptide signal after enrichment (**Figure 5A**). The unbound fraction can then be used to measure relative protein abundance, as confirmed by the analysis in previous sections (**Figure 3–4**). To our knowledge, this is the first example of combined proteomics and phosphoproteomics workflow, using any genetically targeted proximity labeling methods in the mouse brain.

**Figure 5.**
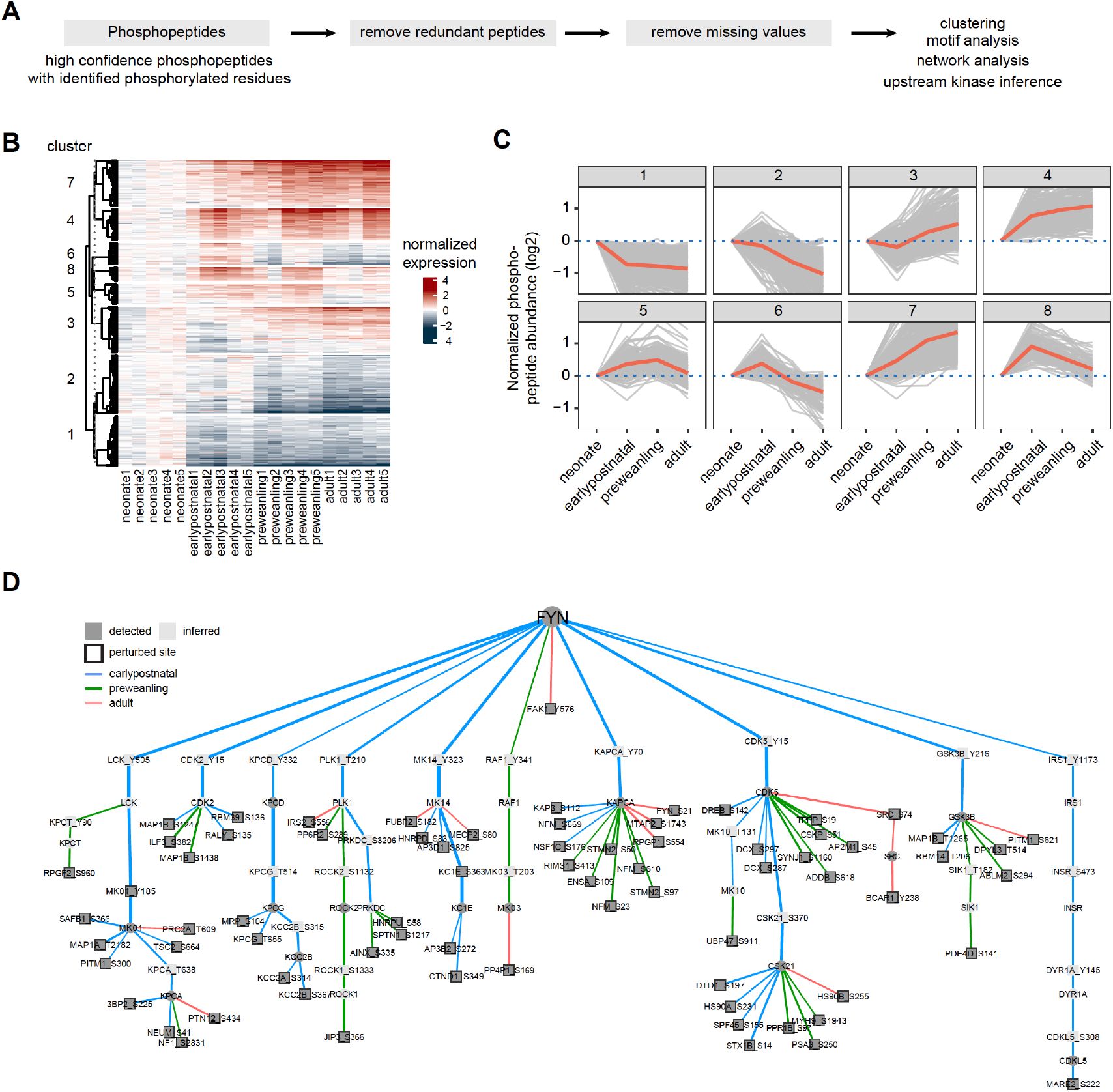
Temporal trajectories of phosphopeptide abundance and kinase-substrate interaction network. **(A)** Phosphopeptide analysis workflow. Raw MS data were analyzed by Proteome Discoverer with false discovery rate (FDR) set at 1% for peptide spectrum matches (PSMs). Only phosphopeptide groups with identified phosphorylated residues were used in the analysis. A unique phosphopeptide is defined by a combination of phosphopeptide and its corresponding protein. Entries without missing abundance values across replicates were used in the timecourse and clustering analysis. **(B)** Heat map showing temporal expression of phosphorylation across development. Peptide clusters were defined using a time-course regression analysis implemented in maSigPro R package. The row dendrogram shows hierarchical clustering of peptide in each cluster. Log2 peptide abundance was normalized to the mean of the neonatal time point. All biological replicates are shown in columns. **(C)** Temporal trajectories of phosphorylation in each peptide cluster. Log2 peptide abundance was normalized to the mean of the neonatal time point. Red lines represent the average expression of all proteins in a cluster. Gray lines represent individual protein trajectories. Neonate (P5), early postnatal (P11), preweanling (P18–20), and adult (P50). **(D)** Kinase-substrate interaction network downstream of *Fyn* kinase. PHOsphorylation Networks for Mass Spectrometry (PHONEMeS) was used to model phosphosites using known prior knowledge network from PhosphoSitePlus database. Nodes, kinase or phosphosite. Node color: dark gray, detected, light gray, inferred by the model. Edges, kinase-substrate interactions. Edge thickness shows the degree of importance inferred for each kinase-substrate interaction. Edge color: blue, green, red for earlypostnatal–neonate, preweanling–neonate, and adult–neonate comparison, respectively. See also **Figure S6**.

Here, we only focused on phosphopeptides that were identified with high confidence (1%FDR), with unambiguous phosphosites, and with quantified proteins. After stringently remove peptides with any missing TMT intensity in any sample, we quantified 2353 unique phosphopeptides that were mapped to 708 proteins (**Data S7**). Batch-effect was corrected with internal reference standard normalization. Global median normalization was used to align sample medians (**Figure S6A**). Many proteins have more than one quantified phosphopeptide, with a median of 2 phosphopeptides per protein (**Figure S6B**). Using the same maSigPro analysis used for proteomic analysis, we divided phosphopeptides into 8 clusters. Clusters 3, 4, and 7 show overall increase in phosphorylation, while clusters 1, 2, 6, and 8 show overall decrease in phosphorylation over development (**Figure 5B–C**). Next, we used motif analysis to look for phosphopeptide sequences that are enriched in our dataset. Motif-x implementation in R (*i.e*., rmotifx R package) (Wagih et al., 2016) was used to identify over-represented motifs against the PhosphoSitePlus as the background database (Hornbeck et al., 2015). We found that the top 3 enriched motifs are sequences that can be phosphorylated by proline-directed kinases (**Figure S6C**). This is consistent with our analysis of the axon guidance pathway and previous reports implicating proline-directed kinases such as *Mapk* family, *Gsk3b*, and *Cdk5*, in neurodevelopment (Igarashi and Okuda, 2019).

To incorporate phosphorylation data in the context of axon development, we aim to build a signaling model that can infer upstream kinases in the context of the data, while taking into account developmental timepoint to identify functional kinase-substrate interactions. We used PHOsphorylation Networks for Mass Spectrometry (PHONEMeS) analysis tool to model our data (Gjerga et al., 2021). PHONEMeS reconstructs phosphorylation signaling networks from known kinase-substrate interaction database (*i.e*., prior knowledge network, PKN), taking into account ‘perturbed’ (differentially detected) phosphosites across time points. With respect to a chosen upstream target kinase, PHONEMeS models phosphorylation signaling propagation from the chosen target node to reach the perturbed phosphosites. Our prior analysis of the *Ntn1-Dcc* pathway points towards *Src* and *Fyn* kinases as key regulators of the axon guidance signaling. To better understand the axon guidance pathway, we focused on *Fyn* as our target kinase that is significantly upregulated in the earlier time windows (**Figure S6D**). For PHONEMeS input, phosphosites that are differentially detected from neonate were considered as perturbed nodes for that time point (*e.g*., early postnatal – neonate). PHONEMeS allows users to configure background network to a species other than human. This is important because exact phosphorylation residues are not readily mapped across species. We used OmnipathR to obtain known mouse kinase-substrate interactions (Türei et al., 2016). We relied on Cytoscape for visualizing PHONEMeS output (**Figure 5D**). The model assigned edge color and thickness based on their significance for that time point, evidenced by the data. From this model, we observed that *Fyn* is important for phosphorylation events in the early postnatal time point, revealed by a greater proportion of blue edges (early postnatal) compared to green (preweanling) and red (adult). We also found that *Fyn* can directly or indirectly regulate proline-directed kinases such as *Mapk1* (MK_01), *Cdk5*, and *Gsk3b*, whose phosphorylation motifs are enriched in our dataset (**Figure S6E**). Thus, the result is in agreement with our pathway-centric analysis (**Figure 4F**) and motif analysis (**Figure S6C**), supporting the model that *Fyn* is one of the key regulators in the axon guidance pathway in the corticostriatal systems. Altogether, phosphosite-centric analysis by PHONEMeS provides a useful roadmap to extract kinase-substrate interactions that are contextualized to the data, as well as inferences about intermediate kinases and their potential roles in the signaling pathways.

### Correlation analysis of phosphopeptide and protein abundance

Phosphorylation plays a crucial role in protein function, including control over enzymatic activity and protein-protein interactions. As protein expression changes, phosphorylation (as a percentage of total protein) may remain steady or change as levels or activity of kinase and phosphatases fluctuate. Our dataset highlights dramatic reshaping of axonal proteome levels across development, which raises the question of how protein activity is altered on top of changing protein levels across development. To address this question, we computed Spearman rank correlations between phosphopeptides and proteins across developmental time (**Figure 6A, Data S8**). P-values associated with correlation coefficients from 2109 peptide-protein pairs were corrected, with 10% FDR as the cutoff for statistical signficance. We found that the majority of phosphopeptide intensities correlate with protein abundance, illustrated by a left-skewed histogram (**Figure 6B**). Out of 309 phosphopeptides with a negative correlation to protein level, 59 decorrelated peptides passed the FDR cutoff (**Figure 6C**). To determine the directionality of changes between protein and peptide abundance across time, we calculated log2 fold change for the corresponding protein between adult and neonate for each peptide. Comparing protein log2 fold change with correlation coefficients, we found 17 decorrelated peptides with negative protein log2 fold change, indicating that phosphorylation increases over time while protein abundance decreases. Analogously, 42 decorrelated peptides with positive protein log2 fold change indicate that phosphorylation decreases over time as protein increases.

**Figure 6.**
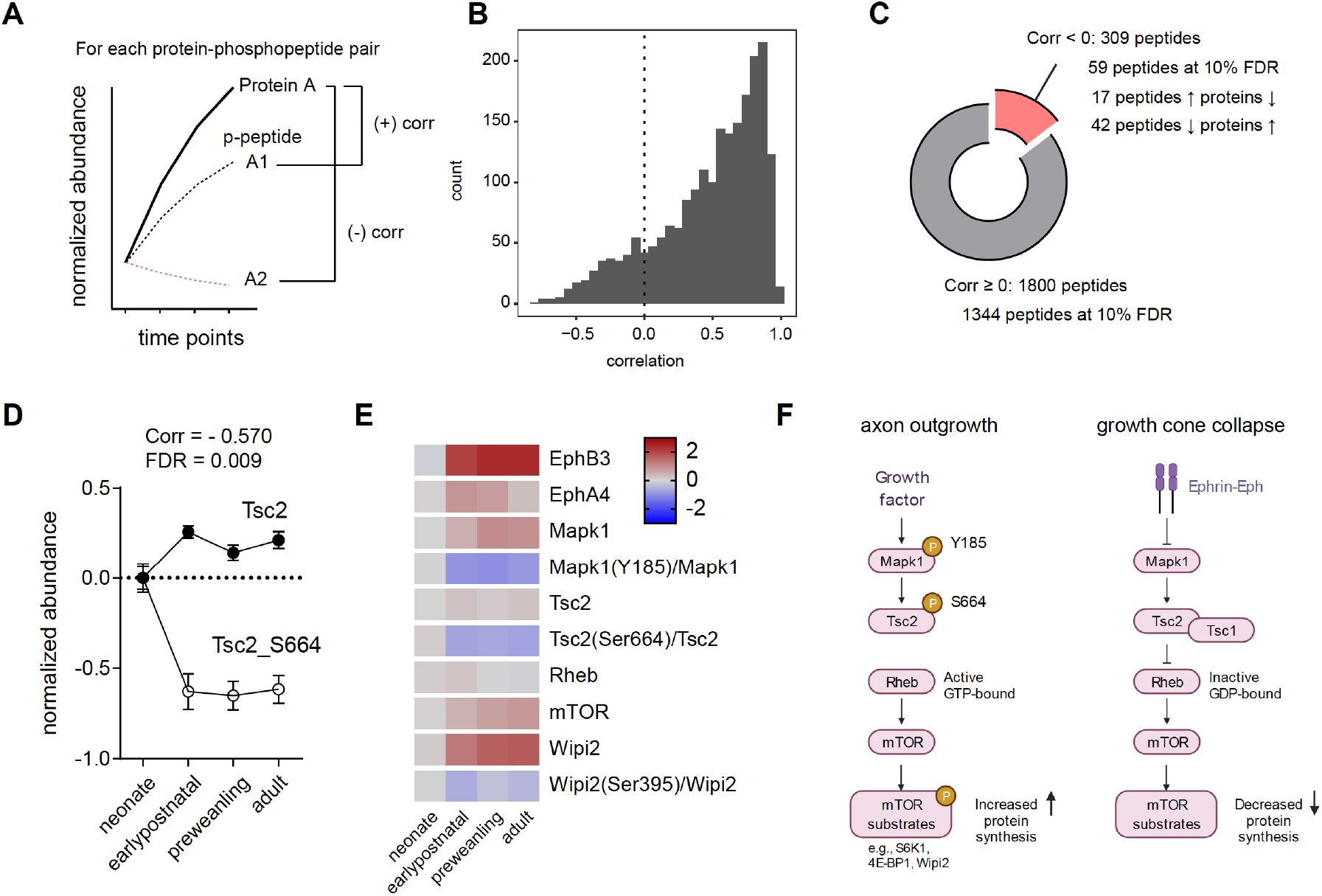
Correlation analysis of phosphorylated peptides and proteins implicates Tsc2 Ser664 in mTOR regulation in corticostriatal axon development. **(A)** Correlation analysis workflow. Spearman rank correlation is calculated for each protein and phosphopeptide pair across time points. One protein may have multiple phosphopeptide-protein pairs, yielding multiple correlation values. False-discovery rate was controlled by the Benjamini-Hochberg approach. **(B)** Distribution of protein-phosphopeptide correlations. Left-skewed histogram (bin width = 1). Most phosphopeptides correlate with their respective protein expression across time. **(C)** Classification of protein-phosphopeptide correlations. Cutoff for correlated peptides, Spearman correlation ≥ 0. 1344 out of 1800 phosphopeptides are positively correlated with proteins by their temporal expression at 10% false discovery rate (FDR). A small proportion of peptides (59 of 309 phosphopeptides) are decorrelated (inversely correlated) at 10% FDR. 42 phosphopeptides decrease in abundance as protein levels increases, while 17 phosphopeptides show the opposite pattern. **(D)** Tsc2 and Tsc2_Ser664 expression. Log2 abundance for Tsc2 (open circles) and Tsc2_Ser664 (solid circles) were normalized to the neonatal time point (Spearman rank correlation: −0.570, FDR = 0.009. n = 5 for each time point. Error bars represent SEM). **(E)** Expression heatmap for proteins and phosphosites in the Tsc2-Rheb-mTOR signaling pathway. Log2 protein expression was normalized to the neonate time point. Log2 phosphopeptide signal was normalized to the corresponding protein abundance and to the neonate time point. **(F)** Model for mTOR-mediated growth cone collapse.

Next, we investigate whether decorrelated phosphosites are important for axonal development. To narrow down the list, we chose to examine decorrelated phosphopeptides on proteins that are associated with disease risk genes (**Data S8**). Among decorrelated peptides, 15 phosphosites are mapped to disease risk genes with 12 out of 15 sites conserved in humans. Particularly, we found Tsc2_Ser664 in our PHONEMeS model under the regulation of Mapk1 (**Figure 5D, Figure S6E**). We found that Tsc2 protein moderately increases throughout development, while Tsc2_Ser664 rapidly drops off after the neonatal period (**Figure 6D**, corr = −0.570, FDR = 0.009). *Tsc2* is one of the key regulators of the mTOR pathway (Ma et al., 2005), and is implicated in autism spectrum disorder. Activation of *Tsc1/Tsc2* complex decreases mTOR activity by inhibiting *Rheb*, a positive mTOR regulator. When *Tsc2* is phosphorylated by *Mapk1* at Ser664, Tsc1/Tsc2 complex is destabilized, allowing *Rheb* to activate mTOR. This interaction between *Mapk1* and Tsc2_Ser664 during the earlier time point is emphasized by the PHONEMeS model. To evaluate this regulation more closely, we generated a simplified model of Tsc2-Rheb-mTOR signaling and plotted relative protein and phosphopeptide abundance normalized to the neonatal time point (**Figure 6E**). In our dataset, relative phosphorylation of *Tsc2* at Ser664 in the corticostriatal axons is highest around P5 and decreases over time. Elevated Tsc2_Ser664 phosphorylation inhibits *Tsc1*/Tsc2 activity, in turn, activating mTOR. We inferred the activity of mTOR by examining its substrate phosphorylation. We detected one mTOR substrate, Wipi2 and Wipi2_Ser395. Consistent with the model, Wipi2_Ser395 phosphorylation is higher in the neonate, supporting the hypothesis that the overall mTOR activity is greater during axon innervation (**Figure 6F**). As neural circuits mature, axonal outgrowth stops and growth cone collapses. A general mechanism for growth cone collapse mediated by ephrin-Eph receptor signaling was studied in the retinal ganglion cells (Nie et al., 2010). The activation of Eph receptor leads to inhibition of *Mapk1*, in turn, allowing *Tsc2* to inhibit *Rheb* and mTOR. In the striatum, it has been reported that the growth cones collapse after P7. As expected, we found *EphA4* and *EphB3* Eph receptor expression increase from P5 to P11, coinciding with reduction in Tsc2_Ser664 and Wipi2_Ser395 (**Figure 6F**). Therefore, our data support the general mechanism that mTOR activity is elevated during axon outgrowth and reduced during growth cone collapse, and that this process is conserved in corticostriatal projections.

In summary, we demonstrate a sensitive proximity labeling workflow involving a tandem enrichment of biotinylated proteins and phosphopeptides. We use a newly generated and validated APEX reporter line to map the axonal proteome in the corticostriatal system and establish an analysis framework to examine the subcellular proteome and phosphoproteome across development, generating broadly applicable resources for studying the proteomic landscape of mammalian neural circuits.

## DISCUSSION

Genetically encoded proximity labeling is an effective way to perform cell-type and subcellular compartment specific proteomics in the mouse brain (Dumrongprechachan et al., 2021; Hobson et al., 2022; Rayaprolu et al., 2021). Here, we generated a new Cre-dependent APEX2 reporter line to capture snapshots of the neuroproteome with cell type specificity, illustrated by several genetic crosses including VGlut2^Cre^, VGAT^Cre^, and Rbp4^Cre^. In comparison with biotin ligase-based reporters (*e.g*., BioID, TurboID), APEX-mediated labeling occurs within 1 hr *ex vivo* in acute brain sections, while *in vivo* BioID/TurboID labeling takes several days up to weeks, via continuous biotin supplementation (Rayaprolu et al., 2021; Spence et al., 2019; Uezu et al., 2016). Therefore, the APEX-based approach is more suitable for biological questions that require short temporal windows, while BioID/TurboID can measure *in vivo* steady-state proteome spanning the duration of biotin administration.

Profiling the axonal proteome or transcriptome is challenging, particularly from complex tissues composed of multiple intermingled cell types. Canonical approaches utilize cell culture models in compartmentalized devices to ensure directional growth (Cagnetta et al., 2018; Chuang et al., 2018), laser capture microdissection of axon terminals (Zivraj et al., 2010), manual dissection of nerve segments (Michaelevski et al., 2010), synaptosome sorting from hippocampal mossy fibers (Apóstolo et al., 2020), or growth cone (GC) sorting from developing brain (Chauhan et al., 2020; Poulopoulos et al., 2019). However, no single approach can flexibly measure the overall axonal proteome across a wide age range, with high efficiency, genetic targeting, and pre-synaptic specificity. In this work, we leverage the early expression of APEX in the transgenic reporter cross to map the proteomic landscape of corticostriatal axons from neonate (P5) to young adult (P50). Enabled by temporal precision of APEX-mediated biotinylation, we demonstrate that APEX-based proximity labeling detects dynamically changing proteome of corticostriatal axons across multiple subcellular compartments including axoplasm, synapses, and mitochondria, without the need for biochemical fractionation, or sample pooling beyond P5.

Furthermore, we have designed a workflow to add additional steps during the sample preparation for enriching phosphorylated peptides from the same samples, as the ex vivo biotinylation yields sufficient materials. Our novel tandem enrichment workflow enables measurement of both protein and phosphopeptide abundance using TMTPro-based multiplexing and quantification. This is crucial for our study, because investigating phosphosite abundance alone without protein abundance can lead to inaccurate interpretation, especially in a system where protein abundance is changing (Wu et al., 2011). While our approach profiles phosphosites on biotinylated proteins, Alice Ting and colleagues recently developed a sequential protein enrichment strategy that combines streptavidin enrichment with functional protein enrichment (*e.g*., phase separation for RNA-binding proteins, IMAC resin for phosphoproteins) to discover novel functions of a protein subclass in a specific subcellular compartment in HEK cells, highlighting the utility of tandem enrichment in the context of APEX based proteomics (Qin et al., 2021).

Using our workflow, we interrogate axonal proteome of the developing corticostriatal system. We obtained sufficient profiling of the overall landscape. Despite stringent removal of proteins with missing measurements, we were able to perform pathway-centric and phosphosite-centric analyses with good coverage. We showcase an important set of systems-level analyses, encompassing a mapping of protein and phosphopeptide temporal trajectories, disease risk gene association, contextualized kinase-substrate interactions, and correlations between protein and phosphopeptide levels across time, enabling interrogation of complex signaling pathways such as the axon guidance. Molecular level mechanisms of axon guidance systems in the brain have been investigated in a limited number of models. Several examples including Netrin-*Dcc* signaling in dopaminergic (Kang et al., 2018; Vosberg et al., 2020), thalamocortical (Castillo-Paterna et al., 2015) and commissural axon projections (Varadarajan et al., 2017), and Ephrin-Eph receptor signaling in retinal ganglion cell projections (Nie et al., 2010), callosal axon navigation (Nishikimi et al., 2011) and hippocampal mossy fibers (Xu and Henkemeyer, 2009). Our pathway-centric analysis reveals that both Netrin-*Dcc* signaling and *EphA* signaling are conserved in corticostriatal projections, demonstrating an analysis framework that synthesizes prior knowledge from different neural circuits. Beyond this example, many signaling pathways remains to be explored and our datasets can be used as a general resource to generate testable models for axon development in the context of neurodevelopmental disorders for further mechanistic studies.

Data generated by this work serve as a general reference for overall corticostriatal afferent development at the proteomic and phosphoproteomic levels. Notably, cortical afferents are classified into pyramidal tract (PT) and intertelencephalic tract (IT) projections (Shepherd, 2013). PT projections innervate the striatum in the embryonic period, while IT projections arise postnatally, around P3–P4 (Nisenbaum et al., 1998; Sheth et al., 1998; Sohur et al., 2014). Both types of cortical projections to the striatum are topographically organized, forming functionally and spatially distinct striatal regions to process and integrate cortical inputs (Hooks et al., 2018). With appropriate Cre-driver lines (*e.g*., Tlx3-Cre_PL56, Sim1-Cre_KJ18 for IT and PT neurons, respectively) (Gerfen et al., 2013) and timing, analogous approaches can be applied to examine the molecular basis of regional input specific proteome from PT or IT projections during early postnatal development. As a proof of principle in this study, we focus on corticostriatal axons during normal development, but axon maturation requires both presynaptic and postsynaptic interactions. The timely formation of striatal circuits including the maturation of striatal neurons depends on proper levels of synaptogenesis and dendritic spinogenesis, regulated by recurrent network activity, neuromodulation, and experience (Kozorovitskiy et al., 2012, 2015). Disruption of axon development by disease associated mutations or absence of appropriate dopamine modulation can result in impaired striatal function (Lieberman et al., 2018; Peixoto et al., 2016). Thus, our work positions the field for genetically targeted proteomics that can be combined with pharmacological, genetic, or engineered effectors to reveal the principles of neural circuit formation between pre- and post-synaptic neurons.

In sum, genetically encoded proteomics tools are rapidly evolving. Together with advances in mass spectrometry, we are starting to obtain sufficient depth of genetically targeted proteome from low microgram samples, with lower rates of missing measurements and with post-translational modification information. With more proteomic datasets of this type, numerous features including cell type, subcellular compartment, developmental trajectory, and phosphorylation can facilitate more accurate theoretical modeling of signaling pathways in the brain during development and in disease contexts. Altogether, methodological and conceptual advancements, along with the datasets generated by this work, provide a new avenue for interrogating the proteomic landscapes of any genetically targeted neural system in the mouse brain.

## MATERIALS AND METHODS

### Mouse strains and genotyping

Animals were handled according to protocols approved by the Northwestern University Animal Care and Use Committee. Weanling and young adult male and female mice were used in this study. Cre transgenic lines used in this study include VGlut2-IRES-Cre (Slc17a6^tm2(Cre)Lowl^, stock. 016963, Jackson Labs), VGAT-IRES-Cre (Slc32a1^tm2(cre)/Lowl^, stock. 016962, Jackson Labs), and Tg(Rbp4-cre)KL100Gsat/Mmucd (stock. 031125-UCD, MMRRC) (Gerfen et al., 2013; Vong et al., 2011). The Cre-dependent APEX2 mouse line (DIO-APEX2.NES-P2A-EGFP) was generated in this study. C57BL/6 mice used for breeding and backcrossing were acquired from Charles River (Wilmington, MA). All mice were group-housed in a humidity-controlled, ambient temperature facility, with standard feeding, 12 hr light-dark cycle, and enrichment procedures. Littermates were randomly assigned to conditions. All animals were genotyped according to the MMRRC strain-specific primers and protocols using GoTaq Green PCR master mix (Cat. No. M712, Promega Corporation, Madison, WI, USA).

## METHOD DETAILS

### Generation of transgenic APEX2 reporter mouse line

DIO-APEX2.NES-P2A-EGFP animals were generated using CRISPR/Cas9 approach at the University of Michigan transgenic core. Mice were housed on ventilated racks or in static microisolator cages with access to food and water with a standard 12 hr light/dark cycle. HDR template was generated by inserting EF1a-DIO-APEX2.NES-P2A-hGH DNA sequence in pROSA26-delDTA between the left and right homology arms. pROSA26-delDTA plasmid was generated by removing the DTA sequence from pROSA26-PA (a gift from Dr. Frank Costantini, Addgene #21271) by cutting with XhoI and SalI, followed by ligation with T4 DNA ligase. crRNA targeting the ROSA26 locus (Chu et al., 2016), 5’-ACTCCAGTCTTTCTAGAAGA-3’ was annealed with tracRNA to form duplex gRNA according to manufacturer’s instructions (Integrated DNA Technologies). The gRNA was combined with enhanced specificity eSpCas9(1.1) protein resuspended in the provided buffer (cat. no. ESPCAS9PRO, Sigma, MO, USA) to form a ribonucleoprotein complex. The pROSA26-delDTA-EF1a-DIO-APEX2.NES-P2A-EGFP-hGH DNA donor plasmid was purified with for microinjection with an endotoxin-free kit and resuspended in RNAse free microinjection buffer (10 mM Tris, 0.25 mM EDTA, pH 7.4). A test injection was performed with the DNA donor to ensure that it did not interfere with the in vitro development of mouse zygotes from the fertilized egg stage through to the blastocyst stage. The DNA donor plasmid was mixed the CRISPR reagent immediately prior to microinjection into mouse zygotes. The final concentration of the reagents was 7.5 ng/ul crRNA + 5 ng/ul tracrRNA (annealed), 25 ng/ul Cas9 protein, 10 ng/ul circular DNA donor plasmid.

Mouse zygote microinjection was carried out as essentially as described (Becker and Jerchow, 2011). Briefly, mixed CRISPR reagents and DNA donor were microinjected into the pronuclei of fertilized eggs at room temperature in hanging drop chambers made of M2 medium (Cat. No. M7167, Sigma) surrounded by mineral oil to prevent evaporation. Microinjection needles were pulled from baked (200°C, 2 hours) 1 mm O.D., 0.75 mm I.D glass capillaries with internal filaments (Cat. No. MTW100F-4, World Precision Instruments) on a Sutter P-87 pipet puller. Microinjection needles were connected to a Femtojet pneumatic microinjector (Eppendorf). Zygote holding pipets (200 um O.D., 30 um I.D.) were fashioned by hand on a microforge from the same glass capillaries and connected to an Eppendorf Cell-Tram Ari. The micromanipulation workstation consisted of a motorized Nikon TE2000–S microscope (Nikon) equipped with differential interference contrast optics and hydraulic hanging joysticks (Narishige). After microinjection of approximately 3 pl of solution the eggs were washed through four drops of 75 ul CO2 equilibrated KSOMaa medium (Zenith Biotech) under mineral oil. Eggs were surgical transferred to pseudopregnant female mice the same day as they were injected. Fertilized eggs were obtained from the mating of superovulated C57BL/6J female mice (The Jackson Laboratory stock no. 000664) or B6SJLF1 female mice (The Jackson Laboratory stock no. 100012) with B6SJLF1 male mice. A total of 308 zygotes were microinjected, 273 zygotes were viable after microinjection and transferred to eleven pseudopregnant recipients. A total of 112 pups were born and screened for the ROSA26 insertion.

### Founder screening, genotyping, and transgene characterization

Mouse tail genomic DNA was extracted in 300 μl DirectPCR lysis solution (Viagen Biotech) supplemented with 0.4 mg/ml Proteinase K (Life technologies) at 55°C overnight. Crude extract was heat inactivated at 85°C for 45min and stored at 4°C. The presence of APEX-transgene was identified using GoTaq Green PCR protocol. Full length transgene and integration sites were evaluated by Q5 PCR protocol. Detailed PCR methods can be found in **Table S1**. Potential off-targets of Cas9 activity were identified via Benchling gRNA design tool. The top eight off-targets with *NGG* PAM sequence were selected for analysis. ~800 bp regions flanking the off-target sites were amplified (**Table S2**). Primers were designed based on sequence retrieved from Mus musculus (assembly GRCm38.p6) reference genome on the NCBI database. 20 μl of PCR product was cleaned up by 2 μl of ExoSAP-IT express reagent for 30 min at 37°C and 2 min at 80°C. Sanger sequencing was used to identify mutations (indels or point mutations). All sequence alignment was performed in SnapGene software.

### Stereotactic injections

Conditional expression of H2B.APEX in corticostriatal projection neurons was achieved by recombinant adeno-associated viral neonatal transduction encoding a double-floxed inverted open reading frame (DIO) of target genes, as described previously (Dumrongprechachan et al., 2021). For neonatal AAV delivery, P3-6 mice were cryoanesthetized and were placed on a cooling pad. 400 nl of AAV was delivered using an UltraMicroPump (World Precision Instruments, Sarasota, FL) by directing the needle +1 mm anterior of bregma, ±0.15 mm from midline, and 0.8-1.0 mm ventral to skin surface. Following the procedure, pups were warmed on a heating pad and returned to home cages, with approved post procedure monitoring. AAVs were diluted to the final titers using Gibco PBS pH 7.4 (AAV1.EF1a.DIO.H2B-APEX-P2A-EGFP titer ~3×10^12^ GC/ml). P35-P70 animals were used for histology, western blots, and proteomics experiments. Cre-negative animals (APEX2 reporter alone) were used as a negative enrichment control group for proteomics, because they do not express APEX transgenes.

### Acute slice preparation and *ex vivo* biotinylation

Animals were deeply anesthetized with isofluorane inhalation. The brain was removed and placed directly into ice-cold carbogenated (95%O_2_/5%CO_2_) artificial cerebrospinal fluid (ACSF; 127 mM NaCl, 25 mM NaHCO_3_, 1.25 mM NaH_2_PO_4_ monobasic monohydrate, 25 mM glucose, 2.5 mM KCl, 1 mM MgCl_2_, and 2 mM CaCl_2_). Tissues were blocked and sliced on Leica VT1000S or VT1200S. For *ex vivo* biotinylation, slices were incubated in carbogenated ACSF with 500 μM biotin phenol (Cat. No., LS-3500, Iris Biotech, Germany) at RT for 1 hour. They were briefly rinsed in ACSF and then transferred into ACSF containing 0.03% H_2_O_2_ for 2 min. The reaction was quenched with ASCF containing 10 mM sodium ascorbate. Prefrontal cortex or striatum were dissected in > 10 ml ice-cold ACSF on sylgard coated petri dish. ACSF was removed and tissues were stored at −80°C until further processing.

### Immunofluorescence staining

Animals were deeply anesthetized with brief isofluorane inhalation and transcardially perfused with 4% paraformaldehyde (PFA) in 0.1 M phosphate buffer saline (PBS). Brains were extracted and post-fixed in 4% PFA overnight. Brain was rinsed three times with PBS. VT1000S was used to generate 40-60 μm sections in PBS for immunohistochemistry or tissue imaging. APEX expressing cells were identified by GFP immunostaining using chicken anti-GFP antibody (1:2,000; Cat. No. AB13970, Abcam, Cambridge, UK, RRID:AB_300798) and APEX localization was confirmed by mouse anti-FLAG antibody (1:1,000; Cat. No. A00187-200, Genscript, NJ, USA, RRID:AB_1720813). Sections were incubated in primary antibody solution overnight at 4°C (0.5% Triton PBS). Tissues were rinsed three times with PBS (5 min each) and incubated in secondary antibody solution at RT for 1hr (PBS 1:500 goat anti-chicken Alexa Fluor 488 (RRID:AB_2534096), and goat biotinylated anti-mouse, Thermo Fisher). For detecting FLAG-tagged APEX, tissues were incubated in 1:500-streptavidin Alexa Fluor 647 (Thermo Fisher) at RT for 1 hr. Tissues were then rinsed 3x (5 min each) with PBS, air dried on Superfrost Plus slides and coverslipped under 10% glycerol-TBS with 2 μg/ml Hoechst 33342 (Thermo Fisher). Sections were then imaged with Olympus VS120 microscope system and/or Leica SP5 confocal microscope.

To verify APEX activity and subcellular localization, tissue sections were incubated in PBS containing 500 μM biotin phenol for 30 min and treated with PBS containing 0.03% H_2_O_2_ for 1min. The reaction was quenched 3x with PBS containing 10 mM NaN_3_ and 10 mM sodium ascorbate. Following biotinylation, tissues were immunostained with 1:2000-chicken anti-GFP, 1:1000-rabbit anti-DARPP32 overnight at 4°C (Cat. No. 2306, Cell Signaling, Danvers, MA, USA, RRID:AB_823479). Tissues were rinsed three times with PBS (5 min each) and incubated in secondary antibody solution at RT for 1hr (PBS 1:500 goat anti-chicken Alexa Fluor 488 (RRID:AB_2534096), goat anti-rabbit Alexa Fluor 594 (RRID:AB_2534079) and streptavidin Alexa Fluor 647, Thermo Fisher).

### Western blot analysis

Total protein was extracted by sonication in 400 μl lysis buffer (1% sodium dodecyl sulfate (SDS), 125 mM triethylammonium bicarbonate (TEAB), 75 mM NaCl, and Halt™ protease and phosphatase inhibitors). Lysates were cleared by centrifugation at 12,000 g for 15 min at 4°C. Supernatant was transferred to a new tube and used for subsequent procedure. Total protein (1 μl of samples were diluted to 100 μl with water) was estimated using microBCA assay according to the manufacturer instructions (Cat. No. 23235, Thermo Fisher).

Protein lysate inputs or flow through fractions were mixed with 6x Laemmli loading buffer and heated to 90-95°C for 10 min. For eluting biotinylated protein off the beads, washed beads were mixed with 20 μl of 2x Laemmli buffer containing 25 mM TEAB, 75 mM NaCl, and 20 mM biotin. Beads were heated to 90-95°C for 10 min. Proteins were separated in 10%, 12%, or 4-20% gradient gels (Cat. No. 4561096, Biorad, CA, USA) and transferred to nitrocellulose membrane (Cat. No. 926-31090, LI-COR, NE, USA). Blots were briefly rinsed with TBS. For detection of biotinylated proteins, blots were incubated in TBST (0.1% Tween-20) containing streptavidin CW800 (1:10,000, Cat. No. 926-32230, LI-COR) for 1 hr at RT. Blots were washed three times with TBST for 10 min each. Total protein was detected using REVERT 700 according to the manufacturer instructions. For other proteins, blots were blocked with 5% milk TBS for 1 hr and probed with primary antibodies prepared in TBST overnight at 4°C (1:2000 for tyrosine hydroxylase, Doublecortin, synaptophysin-I, Psd95, and 1:1000 for DARPP32). Blots were washed three times with TBST 10 min each at RT and probed with 1:10000-secondary antibodies (LI-COR, RRID:AB_10956166, RRID:AB_621843, RRID:AB_10956588, RRID:AB_621842). Blots were scanned using a LI-COR Odyssey CLx scanner. All quantification was performed using LI-COR Image Studio version 5.2. For dot blot analysis, 1 μl (0.2 μg) was dotted on nitrocellulose membrane. Membrane was air dried and probed with 1:20000-streptavidin CW800 and total protein stain REVERT 700.

### Mass spectrometry sample preparation

This protocol was modified from our previous publication (Dumrongprechachan et al., 2021). Total protein was extracted by sonication in lysis buffer (1% SDS, 125 mM TEAB, 75 mM NaCl, and Halt™ protease and phosphatase inhibitors). Lysates were cleared by centrifugation at 12000 g for 15 min at 4°C. Supernatant was used for subsequent procedure. Total protein was estimated using BCA assay according to the manufacturer instructions (Cat. No. 23235, Thermo Fisher).

Brain lysates (300 μg in 250 μl) were reduced with 20 μl 200 mM DTT for 1 hr and alkylated with 60 μl 200 mM IAA in the dark for 45 min at 37°C with shaking. No pooling was needed except for P5 neonatal striatal samples. For the P5 age group, lysates from two animals of the same sex were pooled to make 300 μg. Streptavidin magnetic beads (150 μl for each sample) (Cat. No. 88816, Thermo Fisher) were prewashed with 1ml no-SDS lysis buffer and incubated with 250 μl of reduced and alkylated lysates for 90 min at RT with shaking. Enriched beads were washed twice with 1 ml no-SDS lysis buffer, 1 ml 1 M KCl, and five times with 1 ml 100 mM TEAB buffer. Washed beads were digested with trypsin solution (~4 μg in 150 μl 100 mM TEAB) overnight at 37°C (Cat. No. 90058, Thermo Fisher). Digested supernatant was collected. Beads were rinsed with 50 μl 100 mM TEAB. Supernatant were combined. Trace amount of magnetic beads were removed twice by magnetization and tube changes. 10 μl from each sample was saved for Pierce fluoremetric peptide quantification assay. Remaining peptides were frozen and dried in a vacuum concentrator before TMT labeling. To make reference channel samples, a separate set of P18 Rbp4^Cre+^;APEX^+^ samples were prepared and pooled to make two 900 μg samples (750 μl, reduced with 60 μl 200 mM DTT, and alkylated with 180 μl 200 mM IAA). Each pooled lysates were enriched with 400 μl of streptavidin beads, followed by the same bead washing and tryptic digestion (~8 μg trypsin in 300 μl + 50 μl rinse). Eluted peptides were pooled and dried in a vacuum concentrator.

Dried peptide samples were reconstituted in 20 μl 100 mM TEAB and sonicated for 15 min at RT. Briefly, one set of 0.5 mg TMTPro 16plex reagents was warmed up to room temperature and dissolved in 20 μl Optima LC/MS-grade acetonitrile (ACN). 6 μl of ~59 mM TMTPro reagents was added to 20 μl reconstituted peptide according to the experimental design in **Table S3**. For reference channel sample, peptides were reconstituted in 80 μl. 40 μl was labeled with 12 μl TMTPro 134N. Labeling was performed at RT for 90 min with shaking at 400 rpm. The reaction was quenched by adding 3 μl 5% hydroxylamine/100 mM TEAB at RT for 15 min with shaking (6 μl for reference channel). All samples (for each TMT set, ~29 μl samples x 15 and 24 μl reference) were mixed equally and dried in a vacuum concentrator.

TMT-labeled peptide mixtures were cleaned up by high pH-reverse phase fractionation (Cat. No. 84868, Thermo Fisher). Samples were resuspended and sonicated for 10 min in 300μl buffer A (LC MS water 0.1% TFA). Resin was packed by centrifugation at 5,000 g for 2 min, activated twice with 300 μl ACN, and conditioned twice with 300 μl buffer A. Peptides were loaded once by centrifugation at 3,000 g for 2 min. Column was first washed with 300 μl water, and 300 μl 2% ACN 98% triethylamine (TEA). Peptides were eluted in 300 μl 25% ACN 75% TEA and dried in a vacuum concentrator prior to phosphopeptide enrichment.

Phosphopeptides were enriched using the AssayMAP Agilent Bravo instrument with Fe(III)-NTA IMAC resin according to the manufacturer’s instructions (Agilent, Cat. No. G5496-60085). Peptides were resuspended in 210 μl 80% ACN 0.1% TFA, bath sonicated for 10 minutes, vortexed for 10 minutes, and quickly centrifuged. The resin was primed with 50% ACN 0.1% TFA and equilibrated with equilibration/wash buffer 80% ACN 0.1% TFA. Samples were loaded into the syringes then dispensed into the Flow Through Collection plate. Syringes were washed with the Cartridge Wash Buffer discarded into waste then repeated with Syringe Wash Buffer. Elution buffer was aspirated into the syringes and dispensed into the Eluate Collection plate. Phosphopeptides were eluted in 1% ammonium hydroxide solution and acidified to 1% formic acid for MS analysis. The flow through fraction was collected and fractionated by high pH-reverse phase columns as described above. Peptides were eluted by increasing percentage of acetonitrile in 0.1% triethylamine solution according to the manufacturer instructions (5%, 10%, 12.5%, 15%, 17.5%, 20%, 22.5%, and 25% ACN). All fractions were dried in a vacuum concentrator.

### Mass spectrometry data acquisition and raw data processing

TMT labeled peptides (~1 μg) were resuspended in 2% acetonitrile/0.1% formic acid prior to being loaded onto a heated PepMap RSLC C18 2 μm, 100 angstrom, 75 μm x 50 cm column (ThermoScientific) and eluted over 180 min gradients optimized for each high pH reverse-phase fraction (Dumrongprechachan et al., 2021). Sample eluate was electrosprayed (2,000V) into a Thermo Scientific Orbitrap Eclipse mass spectrometer for analysis. MS1 spectra were acquired at a resolving power of 120,000. MS2 spectra were acquired in the Ion Trap with CID (35%) in centroid mode. Real-time search (max search time = 34 s; max missed cleavages = 1; Xcorr = 1; dCn = 0.1; ppm = 5) was used to select ions for synchronous precursor selection for MS3. MS3 spectra were acquired in the Orbitrap with HCD (60%) with an isolation window = 0.7 m/z and a resolving power of 60,000, and a max injection time of 400 ms. 4 μl (out of 20) of the TMT labeled phosphopeptide enrichments were loaded onto a heated PepMap RSLC C18 2 μm, 100 angstrom, 75 μm x 50 cm column (ThermoScientific) and eluted over a 180 min gradient: 1 min 2% B, 5 min 5% B, 160 min 25% B, 180 min 35% B. Sample eluate was electrosprayed (2,000V) into a Thermo Scientific Orbitrap Eclipse mass spectrometer for analysis. MS1 spectra were acquired at a resolving power of 120,000. MS2 spectra were acquired in the Orbitrap with HCD (38%) in centroid mode with an isolation window = 0.4 m/z, a resolving power of 60,000, and a max injection time of 350 ms.

Raw MS files were processed in Proteome Discoverer version 2.4 (Thermo Scientific, Waltham, MA). MS spectra were searched against the *Mus musculus* Uniprot/SwissProt database. SEQUEST search engine was used (enzyme=trypsin, max. missed cleavage = 4, min. peptide length = 6, precursor tolerance = 10 ppm). Static modifications include carbamidomethyl (C, +57.021 Da), and TMT labeling (N-term and K, +304.207 Da for TMTpro16). Dynamic modifications include oxidation (M, +15.995 Da), Phosphorylation (S, T, Y, +79.966 Da, only for phosphopeptide dataset), acetylation (N-term, +42.011 Da), Met-loss (N-term, −131.040 Da), and Met-loss+Acetyl (N-term, −89.030 Da). PSMs were filtered by the Percolator node (max Delta Cn = 0.05, target FDR (strict) = 0.01, and target FDR (relaxed) = 0.05). Proteins were identified with a minimum of 1 unique peptide and protein-level combined q values < 0.05. Reporter ion quantification was based on corrected S/N values with the following settings: integration tolerance = 20 ppm, method = most confident centroid, co-isolation threshold = 70, and SPS mass matches = 65. PSMs results from Proteome Discoverer were exported for analysis in MSstatsTMT R package (version 1.7.3).

### Evaluation of protein and phosphopeptide abundance and axonal enrichment

Protein summarization and data analysis workflow was adapted from our previous work (Dumrongprechachan et al., 2021). PSMs was exported from Proteome Discoverer and converted into MSstatsTMT-compatible format using PDtoMSstatsTMTFormat function. PSMs were filtered with a co-isolation threshold = 70 and peptide percolator q value < 0.01. For protein quantifications, only unique peptides were used. In addition, only proteins with a minimum of two unique PSMs were quantified (*e.g*., proteins with 1 unique peptide must have at least two PSMs of different charges to be considered for summarization in the MSstatsTMT). Therefore, not all identified proteins were quantified. Protein summarization was performed in MSstatsTMT with the following arguments: method = LogSum, reference normalization = TRUE, imputation = FALSE.

To remove non-specifically enriched proteins, global median normalization was not performed to reflect enrichment differences between APEX+ and APEX-negative control. Instead, we used the moderated t-test implemented in the MSstatsTMT package to make the following comparisons (CTX_H2B – CTX_negative, and STR_P18 – STR_negative). Multiple hypothesis adjustment was performed using the Benjamini Hochberg approach. Proteins were considered positively enriched at 0.5% FDR and 2.5-fold change above Cre-negative control. After removing non-specific enrichment, the extent of axon-enrichment was assessed by moderated t-test comparison between CTX_H2B and STR_P18 samples. For this comparison, protein abundance was normalized at the peptide-level by setting global median normalization = TRUE in the MSstatsTMT function. Batch-effect was corrected using LIMMA removeBatchEffect function, followed by protein-level median normalization. Proteins were considered as axon-enriched at 5% FDR and log2FC (STR_P18 – CTX_H2B) > 0. For gene ontology analysis, DAVID v6.8 (https://david.ncifcrf.gov/) was used with identifier = ‘UNIPROT_ACCESSION’. All quantified proteins were used as background. Axon-enriched and somatic-nuclear protein lists were used as input, and −log10(FDR) was plotted for top GO terms. For rank-based analysis in **Fig. 4SC**, we first obtained the total number of proteins in the dataset, that has been previously annotated with GO-terms related to axons and presynapses, denoted by A. The number of remaining proteins not in A is denoted by B. We calculate the GO annotation rate defined as the cumulative proportions of A and B as a function of rank. The difference in annotation rate (A – B) for each rank reflects the degree of axon-protein enrichment for that rank.

For phosphopeptide analysis, only high confidence phosphopeptides in striatal samples were used. Reference channel normalization and LIMMA batch effect correction were performed as described above. Global median normalization was used to align medians across TMT channels. We considered a phosphopeptide unique if there was only one peptide-protein entry in the dataset. For duplicate entries (e.g., two peptides with the same modified phosphosite, but different additional modifications such as methionine oxidation), an entry with the maximum signal summed across channels was used.

### Time course and clustering analysis with maSigPro

To statistically evaluate whether proteins or phosphopeptides change across postnatal development, maSigPro package (version 1.64.0) was used (Conesa et al., 2006). Only quantified protein with complete cases in the striatal samples were used. For each protein, maSigPro uses the least-square linear regression to determine whether protein or phosphopeptide abundance significantly changes across four time points. P-value of the F-statistics for each ANOVA (i.e., proteins or unique phosphopeptides) were corrected with Benjamini Hochberg method. Protein or phosphopeptides of similar expression across time were grouped by maSigPro clustering approach (n = 8 clusters) using the regression coefficients and hclust method, providing clustering analysis that is based on time course, not abundance alone. Correlation analysis was performed for each phosphopeptide and the corresponding parent protein. Spearman rank correlation was calculated across developmental time. P-values associated with correlation were corrected by Benjamini Hochberg method. The R script can be found in the Kozorovitskiy lab github account (https://github.com/KozorovitskiyLaboratory/STRaxon).

### Overrepresentation analysis (ORA) for genes associated with CNS disorders and Reactome pathway

Hypergeometric test implemented in Webgestal (http://www.webgestalt.org/) was used to test whether protein clusters were overrepresented by any CNS disorders. Lists of risk genes for Autism, Developmental Delay, Epilepsy, ADHD, Schizophrenia, Major Depressive Disorder, Multiple Sclerosis, Parkinson’s Disease, Alzheimer’s disease, and Glioma (**Data S5**) were compiled from recent reports of large GWAS studies of each disorder. Uniprot accessions from protein clusters were used as input. Quantified proteins with complete cases (no missing values) were used as background. Reactome pathway enrichment was performed in Webgestal with the same input and background as described above. Other parameters were default parameters.

### Network analysis for STRING-DB, KEGG axon guidance and phosphosite analysis

For the KEGG pathway analysis, the axon guidance pathway was downloaded from the KEGG website (https://www.genome.jp/pathway/mmu04360). Protein abundance was normalized to the neonate condition. We mapped KEGG genes back to Swissprot and retained entries that were detected in the dataset. We plotted the data as two heatmaps: kinases/phosphatases, and other axon-guidance related proteins. Kinases and phosphatases were identified by cross referencing the protein list with the Uniprot protein kinase family database (https://www.uniprot.org/docs/pkinfam) and the human dephosphorylation database (DEPOD, http://www.depod.bioss.uni-freiburg.de/) (Duan et al., 2015). Interactions related to the Netrin1-Dcc subnetwork and log2FC were entered manually into Cytoscape for visualization. For STRING-DB analysis in **Fig. S5C**, to determine if there is any known protein-protein interactions between autism risk genes and the calmodulin pathway, the following genes detected in our dataset (*Ank2, Ppp1r9b, Adcy1, Calm2, Prkar1b*, and *Prkcg*) were used as the input for the STRING-DB (Szklarczyk et al., 2021). Only nodes with connected edges were plotted.

For motif analysis, we used the rmotifx R package (https://github.com/omarwagih/rmotifx). PhosphoSitePlus (downloaded in Sept 2021) as the background database and the entire phosphopeptides in our dataset (reformatted into 15-amino acid sequences) as the foreground input. Default parameters were used with serine as the central residue, min.seqs = 20, p-value cutoff = 1e-6.

For PHONEMES-ILP analysis, we adapted the code published by the Saez-Rodriguez lab github (https://github.com/saezlab/PHONEMeS-ILP) (Gjerga et al., 2021). We used OmnipathR to download the mouse kinase-substrate interactions and filtered for entries that map to Swissprot. This list was used as the prior knowledge network (PKN). Next, we normalized each phosphopeptide to the neonate condition using LIMMA, resulting in the following comparison groups with log2 fold changes and adjusted p-values (earlypostnatal – neonate, preweanling – neonate, and adult – neonate). We defined phosphosites as ‘perturbed’ if the adjusted p-value for the comparison to the neonate was < 0.01. This threshold was chosen so that there were sufficient number of interactions detected in our dataset that are also found in the PKN. We used IBM cplex implementation of PHONEMES-ILP to find network solution across three time points, using *Fyn* as the target node for n=50 iterations. Interactions that were presented in more than 25 sub-sampling iterations were visualized by Cytoscape.

## Data availability

Raw data generated in this study are available from the corresponding author on reasonable request. Mass spectrometry raw data have been deposited in the PRIDE database (accession number: PXD030864, Reviewer Username: and Password: XWZnPlis).

## Code availability

Analyzed data generated in this study are provided in the Supplemental information. Analysis code is available on Github (https://github.com/KozorovitskiyLaboratory/STRaxon).

## Acknowledgements

The authors are grateful to Lindsey Butler for mouse colony management, Dr. Thomas Bozza for advice during mouse line generation, Dr. Thom Saundersm, Wanda Filipiak, and Galina Gavrilina for preparation of gene-edited mice in the Transgenic Animal Model Core of the University of Michigan’s Biomedical Research Core Facilities, Northwestern Biological Imaging Facility (RRID:SCR_017767) and Dr. Tiffany Schmidt for confocal microscope access, Northwestern High Throughput Analysis laboratory for the microplate reader, and Dr. Anastasia Yocum and her team at A2IDEA, LLC as data analysis consultants. Some schematics were created with BioRender.com.

This work was supported by the NSF CAREER Award 1846234, NIMH R56MH113923, NINDS R01NS107539, NIMH R01MH117111, the Beckman Young Investigator Award, Searle Scholar Award, Rita Allen Foundation Scholar Award, and Sloan Research Fellowship (all Y.K.), and NIMH R01MH118497 (M.L.M.). V.D. is a predoctoral fellow of the American Heart Association (19PRE34380056) and an affiliate fellow of the NIH 2T32GM15538.

## Author contributions

V.D., M.L.M., and Y.K. designed the study. V.D. carried out most experiments and analyses in the study. V.D. and L.B. generated and validated the APEX2 mouse line. M.L.M. and R.B.S. performed MS sample preparation, data acquisition and analyses. V.D., M.L.M., and Y.K. wrote the manuscript, with feedback from all authors.

## Supplementary Information

Figures S1–6

Tables S1–3

Data S1–8

## REFERENCES

Abrahams, B.S., Arking, D.E., Campbell, D.B., Mefford, H.C., Morrow, E.M., Weiss, L.A., Menashe, I., Wadkins, T., Banerjee-Basu, S., and Packer, A. (2013). SFARI Gene 2.0: a community-driven knowledgebase for the autism spectrum disorders (ASDs). Mol Autism 4, 36.

Alvarez-Castelao, B., Schanzenbächer, C.T., Hanus, C., Glock, C., tom Dieck, S., Dörrbaum, A.R., Bartnik, I., Nassim-Assir, B., Ciirdaeva, E., Mueller, A., et al. (2017). Cell-type-specific metabolic labeling of nascent proteomes in vivo. Nat Biotechnol 35, 1196–1201.

Apóstolo, N., Smukowski, S.N., Vanderlinden, J., Condomitti, G., Rybakin, V., ten Bos, J., Trobiani, L., Portegies, S., Vennekens, K.M., Gounko, N.V., et al. (2020). Synapse type-specific proteomic dissection identifies IgSF8 as a hippocampal CA3 microcircuit organizer. Nat Commun 11, 5171.

Becker, K., and Jerchow, B. (2011). Generation of transgenic mice by pronuclear microinjection. In Advanced Protocols for Animal Transgenesis, (Springer), pp. 99–115.

Cagnetta, R., Frese, C.K., Shigeoka, T., Krijgsveld, J., and Holt, C.E. (2018). Rapid Cue-Specific Remodeling of the Nascent Axonal Proteome. Neuron 99, 29–46.e4.

Carlyle, B.C., Kitchen, R.R., Kanyo, J.E., Voss, E.Z., Pletikos, M., Sousa, A.M.M., Lam, T.T., Gerstein, M.B., Sestan, N., and Nairn, A.C. (2017). A multiregional proteomic survey of the postnatal human brain. Nat Neurosci 20, 1787–1795.

Castillo-Paterna, M., Moreno-Juan, V., Filipchuk, A., Rodríguez-Malmierca, L., Susín, R., and López-Bendito, G. (2015). DCC functions as an accelerator of thalamocortical axonal growth downstream of spontaneous thalamic activity. EMBO Rep 16, 851–862.

Chang, D., Nalls, M.A., Hallgrímsdóttir, I.B., Hunkapiller, J., van der Brug, M., Cai, F., International Parkinson’s Disease Genomics Consortium, 23andMe Research Team, Kerchner, G.A., Ayalon, G., et al. (2017). A meta-analysis of genome-wide association studies identifies 17 new Parkinson’s disease risk loci. Nat Genet 49, 1511–1516.

Chauhan, M.Z., Arcuri, J., Park, K.K., Zafar, M.K., Fatmi, R., Hackam, A.S., Yin, Y., Benowitz, L., Goldberg, J.L., Samarah, M., et al. (2020). Multi-Omic Analyses of Growth Cones at Different Developmental Stages Provides Insight into Pathways in Adult Neuroregeneration. IScience 23, 100836.

Chen, C.-L., Hu, Y., Udeshi, N.D., Lau, T.Y., Wirtz-Peitz, F., He, L., Ting, A.Y., Carr, S.A., and Perrimon, N. (2015). Proteomic mapping in live Drosophila tissues using an engineered ascorbate peroxidase. PNAS 112, 12093–12098.

Chu, V.T., Weber, T., Graf, R., Sommermann, T., Petsch, K., Sack, U., Volchkov, P., Rajewsky, K., and Kühn, R. (2016). Efficient generation of Rosa26 knock-in mice using CRISPR/Cas9 in C57BL/6 zygotes. BMC Biotechnology 16, 4.

Chuang, C.-F., King, C.-E., Ho, B.-W., Chien, K.-Y., and Chang, Y.-C. (2018). Unbiased Proteomic Study of the Axons of Cultured Rat Cortical Neurons. J. Proteome Res. 17, 1953–1966.

Colantuoni, C., Lipska, B.K., Ye, T., Hyde, T.M., Tao, R., Leek, J.T., Colantuoni, E.A., Elkahloun, A.G., Herman, M.M., Weinberger, D.R., et al. (2011). Temporal dynamics and genetic control of transcription in the human prefrontal cortex. Nature 478, 519–523.

Conesa, A., Nueda, M.J., Ferrer, A., and Talón, M. (2006). maSigPro: a method to identify significantly differential expression profiles in time-course microarray experiments. Bioinformatics 22, 1096–1102.

Consortium, T.S.W.G. of the P.G., Ripke, S., Walters, J.T., and O’Donovan, M.C. (2020). Mapping genomic loci prioritises genes and implicates synaptic biology in schizophrenia. MedRxiv, https://doi.org/10.1101/2020.09.12.20192922

Costa-Mattioli, M., Sossin, W.S., Klann, E., and Sonenberg, N. (2009). Translational Control of Long-Lasting Synaptic Plasticity and Memory. Neuron 61, 10–26.

Dani, J.W., Armstrong, D.M., and Benowitz, L.I. (1991). Mapping the development of the rat brain by GAP-43 immunocytochemistry. Neuroscience 40, 277–287.

Deciphering Developmental Disorders Study (2017). Prevalence and architecture of de novo mutations in developmental disorders. Nature 542, 433–438.

Demontis, D., Walters, R.K., Martin, J., Mattheisen, M., Als, T.D., Agerbo, E., Baldursson, G., Belliveau, R., Bybjerg-Grauholm, J., Bækvad-Hansen, M., et al. (2019). Discovery of the first genome-wide significant risk loci for attention deficit/hyperactivity disorder. Nat Genet 51, 63–75.

Duan, G., Li, X., and Köhn, M. (2015). The human DEPhOsphorylation database DEPOD: a 2015 update. Nucleic Acids Res 43, D531–535.

Dumrongprechachan, V., Salisbury, R.B., Soto, G., Kumar, M., MacDonald, M.L., and Kozorovitskiy, Y. (2021). Cell-type and subcellular compartment-specific APEX2 proximity labeling reveals activity-dependent nuclear proteome dynamics in the striatum. Nat Commun 12, 4855.

Fu, J.M., Satterstrom, F.K., Peng, M., Brand, H., Collins, R.L., Dong, S., Klei, L., Stevens, C.R., Cusick, C., Babadi, M., et al. (2021). Rare coding variation illuminates the allelic architecture, risk genes, cellular expression patterns, and phenotypic context of autism. medRxiv, https://doi.org/10.1101/2021.12.20.21267194

Gerfen, C.R., Paletzki, R., and Heintz, N. (2013). GENSAT BAC Cre-Recombinase Driver Lines to Study the Functional Organization of Cerebral Cortical and Basal Ganglia Circuits. Neuron 80, 1368–1383.

Gjerga, E., Dugourd, A., Tobalina, L., Sousa, A., and Saez-Rodriguez, J. (2021). PHONEMeS: Efficient Modeling of Signaling Networks Derived from Large-Scale Mass Spectrometry Data. J. Proteome Res. 20, 2138–2144.

Glock, C., Biever, A., Tushev, G., Nassim-Assir, B., Kao, A., Bartnik, I., Tom Dieck, S., and Schuman, E.M. (2021). The translatome of neuronal cell bodies, dendrites, and axons. Proc Natl Acad Sci U S A 118, e2113929118.

Gonzalez-Lozano, M.A., Klemmer, P., Gebuis, T., Hassan, C., van Nierop, P., van Kesteren, R.E., Smit, A.B., and Li, K.W. (2016). Dynamics of the mouse brain cortical synaptic proteome during postnatal brain development. Sci Rep 6, 35456.

Hafner, A.-S., Donlin-Asp, P.G., Leitch, B., Herzog, E., and Schuman, E.M. (2019). Local protein synthesis is a ubiquitous feature of neuronal pre- and postsynaptic compartments. Science 364, eaau3644.

Heyne, H.O., Singh, T., Stamberger, H., Abou Jamra, R., Caglayan, H., Craiu, D., De Jonghe, P., Guerrini, R., Helbig, K.L., Koeleman, B.P.C., et al. (2018). De novo variants in neurodevelopmental disorders with epilepsy. Nat Genet 50, 1048–1053.

Hobson, B.D., Choi, S.J., Mosharov, E.V., Soni, R.K., Sulzer, D., and Sims, P.A. (2022). Subcellular proteomics of dopamine neurons in the mouse brain. ELife 11, e70921.

Hooks, B.M., Papale, A.E., Paletzki, R.F., Feroze, M.W., Eastwood, B.S., Couey, J.J., Winnubst, J., Chandrashekar, J., and Gerfen, C.R. (2018). Topographic precision in sensory and motor corticostriatal projections varies across cell type and cortical area. Nat Commun 9, 3549.

Hornbeck, P.V., Zhang, B., Murray, B., Kornhauser, J.M., Latham, V., and Skrzypek, E. (2015). PhosphoSitePlus, 2014: mutations, PTMs and recalibrations. Nucleic Acids Res 43, D512–520.

Huang, D.W., Sherman, B.T., and Lempicki, R.A. (2009). Systematic and integrative analysis of large gene lists using DAVID bioinformatics resources. Nature Protocols 4, 44–57.

Huang, T., Choi, M., Tzouros, M., Golling, S., Pandya, N.J., Banfai, B., Dunkley, T., and Vitek, O. (2020). MSstatsTMT: Statistical Detection of Differentially Abundant Proteins in Experiments with Isobaric Labeling and Multiple Mixtures. Mol Cell Proteomics 19, 1706–1723.

Hung, V., Udeshi, N.D., Lam, S.S., Loh, K.H., Cox, K.J., Pedram, K., Carr, S.A., and Ting, A.Y. (2016). Spatially resolved proteomic mapping in living cells with the engineered peroxidase APEX2. Nat Protoc 11, 456–475.

Hunnicutt, B.J., Jongbloets, B.C., Birdsong, W.T., Gertz, K.J., Zhong, H., and Mao, T. (2016). A comprehensive excitatory input map of the striatum reveals novel functional organization. ELife 5, e19103.

Igarashi, M., and Okuda, S. (2019). Evolutionary analysis of proline-directed phosphorylation sites in the mammalian growth cone identified using phosphoproteomics. Molecular Brain 12, 53.

International Multiple Sclerosis Genetics Consortium (2019). Multiple sclerosis genomic map implicates peripheral immune cells and microglia in susceptibility. Science 365, eaav7188.

Jassal, B., Matthews, L., Viteri, G., Gong, C., Lorente, P., Fabregat, A., Sidiropoulos, K., Cook, J., Gillespie, M., Haw, R., et al. (2020). The reactome pathway knowledgebase. Nucleic Acids Res 48, D498–D503.

Kanehisa, M., Furumichi, M., Tanabe, M., Sato, Y., and Morishima, K. (2017). KEGG: new perspectives on genomes, pathways, diseases and drugs. Nucleic Acids Res 45, D353–D361.

Kang, D.-S., Yang, Y.R., Lee, C., Park, B., Park, K.I., Seo, J.K., Seo, Y.K., Cho, H., Lucio, C., and Suh, P.-G. (2018). Netrin-1/DCC-mediated PLCγ1 activation is required for axon guidance and brain structure development. EMBO Rep 19, e46250.

Keller, D., Erö, C., and Markram, H. (2018). Cell Densities in the Mouse Brain: A Systematic Review. Frontiers in Neuroanatomy 12, 83.

Kozorovitskiy, Y., Saunders, A., Johnson, C.A., Lowell, B.B., and Sabatini, B.L. (2012). Recurrent network activity drives striatal synaptogenesis. Nature 485, 646–650.

Kozorovitskiy, Y., Peixoto, R., Wang, W., Saunders, A., and Sabatini, B.L. (2015). Neuromodulation of excitatory synaptogenesis in striatal development. ELife 4, e10111.

Krajeski, R.N., Macey-Dare, A., van Heusden, F., Ebrahimjee, F., and Ellender, T.J. (2019). Dynamic postnatal development of the cellular and circuit properties of striatal D1 and D2 spiny projection neurons. The Journal of Physiology 597, 5265–5293.

Krogager, T.P., Ernst, R.J., Elliott, T.S., Calo, L., Beránek, V., Ciabatti, E., Spillantini, M.G., Tripodi, M., Hastings, M.H., and Chin, J.W. (2018). Labeling and identifying cell-specific proteomes in the mouse brain. Nat Biotechnol 36, 156–159.

Kuo, H.-Y., and Liu, F.-C. (2019). Synaptic Wiring of Corticostriatal Circuits in Basal Ganglia: Insights into the Pathogenesis of Neuropsychiatric Disorders. ENeuro 6.

Lam, S.S., Martell, J.D., Kamer, K.J., Deerinck, T.J., Ellisman, M.H., Mootha, V.K., and Ting, A.Y. (2015). Directed evolution of APEX2 for electron microscopy and proximity labeling. Nat Methods 12, 51–54.

Lieberman, O.J., McGuirt, A.F., Mosharov, E.V., Pigulevskiy, I., Hobson, B.D., Choi, S., Frier, M. D., Santini, E., Borgkvist, A., and Sulzer, D. (2018). Dopamine Triggers the Maturation of Striatal Spiny Projection Neuron Excitability during a Critical Period. Neuron 99, 540–554.e4.

Lin, A.C., and Holt, C.E. (2007). Local translation and directional steering in axons. EMBO J 26, 3729–3736.

Lobingier, B.T., Hüttenhain, R., Eichel, K., Miller, K.B., Ting, A.Y., Zastrow, M. von, and Krogan, N. J. (2017). An Approach to Spatiotemporally Resolve Protein Interaction Networks in Living Cells. Cell 169, 350–360.e12.

Ma, L., Chen, Z., Erdjument-Bromage, H., Tempst, P., and Pandolfi, P.P. (2005). Phosphorylation and functional inactivation of TSC2 by Erk implications for tuberous sclerosis and cancer pathogenesis. Cell 121, 179–193.

Meriane, M., Tcherkezian, J., Webber, C.A., Danek, E.I., Triki, I., McFarlane, S., Bloch-Gallego, E., and Lamarche-Vane, N. (2004). Phosphorylation of DCC by Fyn mediates Netrin-1 signaling in growth cone guidance. J Cell Biol 167, 687–698.

Michaelevski, I., Medzihradszky, K.F., Lynn, A., Burlingame, A.L., and Fainzilber, M. (2010). Axonal transport proteomics reveals mobilization of translation machinery to the lesion site in injured sciatic nerve. Mol Cell Proteomics 9, 976–987.

Miyamoto, H., Tatsukawa, T., Shimohata, A., Yamagata, T., Suzuki, T., Amano, K., Mazaki, E., Raveau, M., Ogiwara, I., Oba-Asaka, A., et al. (2019). Impaired cortico-striatal excitatory transmission triggers epilepsy. Nat Commun 10, 1917.

Nie, D., Di Nardo, A., Han, J.M., Baharanyi, H., Kramvis, I., Huynh, T., Dabora, S., Codeluppi, S., Pandolfi, P.P., Pasquale, E.B., et al. (2010). Tsc2-Rheb signaling regulates EphA-mediated axon guidance. Nat Neurosci 13, 163–172.

Nisenbaum, L.K., Webster, S.M., Chang, S.L., McQueeney, K.D., and LoTurco, J.J. (1998). Early patterning of prelimbic cortical axons to the striatal patch compartment in the neonatal mouse. Dev Neurosci 20, 113–124.

Nishikimi, M., Oishi, K., Tabata, H., Torii, K., and Nakajima, K. (2011). Segregation and Pathfinding of Callosal Axons through EphA3 Signaling. J. Neurosci. 31, 16251–16260.

Paek, J., Kalocsay, M., Staus, D.P., Wingler, L., Pascolutti, R., Paulo, J.A., Gygi, S.P., and Kruse, A.C. (2017). Multidimensional Tracking of GPCR Signaling via Peroxidase-Catalyzed Proximity Labeling. Cell 169, 338–349.e11.

Peixoto, R.T., Wang, W., Croney, D.M., Kozorovitskiy, Y., and Sabatini, B.L. (2016). Early hyperactivity and precocious maturation of corticostriatal circuits in Shank3B-/-mice. Nat Neurosci 19, 716–724.

Poulopoulos, A., Murphy, A.J., Ozkan, A., Davis, P., Hatch, J., Kirchner, R., and Macklis, J.D. (2019). Subcellular transcriptomes and proteomes of developing axon projections in the cerebral cortex. Nature 565, 356–360.

Qin, W., Myers, S.A., Carey, D.K., Carr, S.A., and Ting, A.Y. (2021). Spatiotemporally-resolved mapping of RNA binding proteins via functional proximity labeling reveals a mitochondrial mRNA anchor promoting stress recovery. Nat Commun 12, 4980.

Rayaprolu, S., Bitarafan, S., Betarbet, R., Sunna, S., Cheng, L., Xiao, H., Bagchi, P., Duong, D.M., Nelson, R., Goettemoeller, A.M., et al. (2021). Cell type-specific biotin labeling in vivo resolves regional neuronal proteomic differences in mouse brain. BioRxiv, https://doi.org/10.1101/2021.08.03.454921

Reinke, A.W., Mak, R., Troemel, E.R., and Bennett, E.J. In vivo mapping of tissue- and subcellular-specific proteomes in Caenorhabditis elegans. Science Advances 3, e1602426.

Rice, T., Lachance, D.H., Molinaro, A.M., Eckel-Passow, J.E., Walsh, K.M., Barnholtz-Sloan, J., Ostrom, Q.T., Francis, S.S., Wiemels, J., Jenkins, R.B., et al. (2016). Understanding inherited genetic risk of adult glioma - a review. Neurooncol Pract 3, 10–16.

Saunders, A., Macosko, E.Z., Wysoker, A., Goldman, M., Krienen, F.M., de Rivera, H., Bien, E., Baum, M., Bortolin, L., Wang, S., et al. (2018). Molecular Diversity and Specializations among the Cells of the Adult Mouse Brain. Cell 174, 1015–1030.e16.

Schanzenbächer, C.T., Langer, J.D., and Schuman, E.M. (2018). Time- and polarity-dependent proteomic changes associated with homeostatic scaling at central synapses. ELife 7, e33322.

Shekarabi, M., Moore, S.W., Tritsch, N.X., Morris, S.J., Bouchard, J.-F., and Kennedy, T.E. (2005). Deleted in Colorectal Cancer Binding Netrin-1 Mediates Cell Substrate Adhesion and Recruits Cdc42, Rac1, Pak1, and N-WASP into an Intracellular Signaling Complex That Promotes Growth Cone Expansion. J. Neurosci. 25, 3132–3141.

Shepherd, G.M.G. (2013). Corticostriatal connectivity and its role in disease. Nat Rev Neurosci 14, 278–291.

Sheth, A.N., McKee, M.L., and Bhide, P.G. (1998). The Sequence of Formation and Development of Corticostriate Connections in Mice. DNE 20, 98–112.

Sohur, U.S., Padmanabhan, H.K., Kotchetkov, I.S., Menezes, J.R.L., and Macklis, J.D. (2014). Anatomic and Molecular Development of Corticostriatal Projection Neurons in Mice. Cerebral Cortex 24, 293–303.

Spence, E.F., Dube, S., Uezu, A., Locke, M., Soderblom, E.J., and Soderling, S.H. (2019). In vivo proximity proteomics of nascent synapses reveals a novel regulator of cytoskeleton-mediated synaptic maturation. Nat Commun 10, 386.

Stahl, E.A., Breen, G., Forstner, A.J., McQuillin, A., Ripke, S., Trubetskoy, V., Mattheisen, M., Wang, Y., Coleman, J.R.I., Gaspar, H.A., et al. (2019). Genome-wide association study identifies 30 loci associated with bipolar disorder. Nat Genet 51, 793–803.

Szklarczyk, D., Gable, A.L., Nastou, K.C., Lyon, D., Kirsch, R., Pyysalo, S., Doncheva, N.T., Legeay, M., Fang, T., Bork, P., et al. (2021). The STRING database in 2021: customizable protein-protein networks, and functional characterization of user-uploaded gene/measurement sets. Nucleic Acids Res 49, D605–D612.

Türei, D., Korcsmáros, T., and Saez-Rodriguez, J. (2016). OmniPath: guidelines and gateway for literature-curated signaling pathway resources. Nat Methods 13, 966–967.

Uezu, A., Kanak, D.J., Bradshaw, T.W.A., Soderblom, E.J., Catavero, C.M., Burette, A.C., Weinberg, R.J., and Soderling, S.H. (2016). Identification of an Elaborate Complex Mediating Postsynaptic Inhibition. Science 353, 1123–1129.

Varadarajan, S.G., Kong, J.H., Phan, K.D., Kao, T.-J., Panaitof, S.C., Cardin, J., Eltzschig, H., Kania, A., Novitch, B.G., and Butler, S.J. (2017). Netrin1 Produced by Neural Progenitors, Not Floor Plate Cells, Is Required for Axon Guidance in the Spinal Cord. Neuron 94, 790–799.e3.

Vong, L., Ye, C., Yang, Z., Choi, B., Chua, S., and Lowell, B.B. (2011). Leptin Action on GABAergic Neurons Prevents Obesity and Reduces Inhibitory Tone to POMC Neurons. Neuron 71, 142–154.

Vosberg, D.E., Leyton, M., and Flores, C. (2020). The Netrin-1/DCC guidance system: dopamine pathway maturation and psychiatric disorders emerging in adolescence. Mol Psychiatry 25, 297–307.

Wagih, O., Sugiyama, N., Ishihama, Y., and Beltrao, P. (2016). Uncovering Phosphorylation-Based Specificities through Functional Interaction Networks. Mol Cell Proteomics 15, 236–245.

Wang, J., Vasaikar, S., Shi, Z., Greer, M., and Zhang, B. (2017). WebGestalt 2017: a more comprehensive, powerful, flexible and interactive gene set enrichment analysis toolkit. Nucleic Acids Research 45, W130–W137.

Werling, D.M., Pochareddy, S., Choi, J., An, J.-Y., Sheppard, B., Peng, M., Li, Z., Dastmalchi, C., Santpere, G., Sousa, A.M.M., et al. (2020). Whole-Genome and RNA Sequencing Reveal Variation and Transcriptomic Coordination in the Developing Human Prefrontal Cortex. Cell Rep 31, 107489.

Wightman, D.P., Jansen, I.E., Savage, J.E., Shadrin, A.A., Bahrami, S., Holland, D., Rongve, A., Børte, S., Winsvold, B.S., Drange, O.K., et al. (2021). A genome-wide association study with 1,126,563 individuals identifies new risk loci for Alzheimer’s disease. Nat Genet 53, 1276–1282.

Winnubst, J., Bas, E., Ferreira, T.A., Wu, Z., Economo, M.N., Edson, P., Arthur, B.J., Bruns, C., Rokicki, K., Schauder, D., et al. (2019). Reconstruction of 1,000 Projection Neurons Reveals New Cell Types and Organization of Long-Range Connectivity in the Mouse Brain. Cell 179, 268–281.e13.

Wray, N.R., Ripke, S., Mattheisen, M., Trzaskowski, M., Byrne, E.M., Abdellaoui, A., Adams, M.J., Agerbo, E., Air, T.M., Andlauer, T.M.F., et al. (2018). Genome-wide association analyses identify 44 risk variants and refine the genetic architecture of major depression. Nat Genet 50, 668–681.

Wu, R., Dephoure, N., Haas, W., Huttlin, E.L., Zhai, B., Sowa, M.E., and Gygi, S.P. (2011). Correct Interpretation of Comprehensive Phosphorylation Dynamics Requires Normalization by Protein Expression Changes. Mol Cell Proteomics 10, M111.009654.

Xu, N.-J., and Henkemeyer, M. (2009). Ephrin-B3 reverse signaling through Grb4 and cytoskeletal regulators mediates axon pruning. Nat Neurosci 12, 268–276.

Zeisel, A., Hochgerner, H., Lönnerberg, P., Johnsson, A., Memic, F., van der Zwan, J., Häring, M., Braun, E., Borm, L.E., La Manno, G., et al. (2018). Molecular Architecture of the Mouse Nervous System. Cell 174, 999–1014.e22.

Zivraj, K.H., Tung, Y.C.L., Piper, M., Gumy, L., Fawcett, J.W., Yeo, G.S.H., and Holt, C.E. (2010). Subcellular Profiling Reveals Distinct and Developmentally Regulated Repertoire of Growth Cone mRNAs. J. Neurosci. 30, 15464–15478.

