## Supplemental Information for "Dynamic proteomic and phosphoproteomic atlas of corticostriatal axon neurodevelopment"

**Table S1. Primers and PCR protocol for APEX reporter line genotyping.**

| Primer name | Sequence (5’ -> 3’) |
|---|---|
| **Q5 amplification of the APEX transgene** | |
| R26R2 | TCTGTGGGAAGTCTTGTCCCTCC |
| R26F1(AAG) | CCAAAGTCGCTCTGAGTTGTTATCAGTAAG |
| | 1x Q5 reaction buffer<br>0.5μM forward and reverse primers<br>200μM mixed dNTPs<br>0.02 U/μl Q5 polymerase<br>1M Betaine (PCR additives from Sigma)<br>1μL heat-inactivated crude lysate (in DirectPCR Lysis solution)<br>Nuclease-free water up to 25μL<br><br>Thermocycling parameters (optimized using gradient PCR):<br>98°C 2min<br>98°C 30sec \| x 30 cycles<br>63°C 30sec \|<br>72°C 3min \|<br>72°C 10min<br>4°C hold<br><br>317bp for WT<br>4156bp for APEX-positive mutant |
| **Q5 amplification of HA-R integration site** | |
| R26F5 | GAAGTTATGTTCTGAGACCATTCTCAGTGGC |
| APEX-N4 | ACAGCGATGTCAAGACCGTTGTTAGC |
| | 1x Q5 reaction buffer<br>0.5μM forward and reverse primers<br>200μM mixed dNTPs<br>0.02 U/μl Q5 polymerase<br>0.5μL heat-inactivated crude lysate (in DirectPCR Lysis solution)<br>Nuclease-free water up to 25μL<br><br>Thermocycling parameters (optimized using gradient PCR):<br>98°C 2min<br>98°C 30sec \| x 40 cycles<br>63°C 30sec \|<br>72°C 3min \|<br>72°C 5min<br>4°C hold<br><br>Expected 5409bp |
| **Q5 amplification of HA-L integration site** | |

| | |
|---|---|
| R26R5 | AGCCTCGGCTAGGTAGGGGATCG |
| EGFP-C | CATGGTCCTGCTGGAGTTCGTG |
| | 1x Q5 reaction buffer<br>0.5µM forward and reverse primers<br>200µM mixed dNTPs<br>0.02 U/µl Q5 polymerase<br>1M Betaine (PCR additives from Sigma)<br>0.5µL heat-inactivated crude lysate (in DirectPCR Lysis solution)<br>Nuclease-free water up to 25µL<br><br>Thermocycling parameters (optimized using gradient PCR):<br>98°C 2min<br>98°C 30sec \| x 40 cycles<br>72.5°C 30sec \|<br>72°C 2min \|<br>72°C 5min<br>4°C hold<br>Expected 2586bp |
| **GoTaq Green PCR for APEX transgene** | |
| R26R3 | TCATACTGTAGTAAGGATCTCAAGCAGG |
| APEX-N4 | ACAGCGATGTCAAGACCGTTGTTAGC |
| | 1x GoTaq Green master mix (Promega)<br>0.1µM forward and reverse primers<br>1µL heat-inactivated crude lysate (in DirectPCR Lysis solution)<br>Nuclease-free water up to 25µL<br><br>Thermocycling parameters (optimized using gradient PCR):<br>95°C 2min<br>95°C 30sec \| x 35 cycles<br>62°C 1min \|<br>72°C 1:40min \|<br>72°C 5min<br>4°C hold<br>Expected 1249bp |

**Table S2. Off-target primers and PCR protocol.**
All off-target PCR was performed using the same protocol.

| **GoTaq Green PCR for potential off-targets** | |
|---|---|
| | 1x GoTaq Green master mix (Promega)<br>0.5μM forward and reverse primers<br>1μL heat-inactivated crude lysate (in DirectPCR Lysis solution)<br>Nuclease-free water up to 50μL<br><br>Thermocycling parameters (optimized using gradient PCR):<br>95°C 5min<br>95°C 30sec \| x 30 cycles<br>55°C 30sec \|<br>72°C 1min \|<br>72°C 5min<br>4°C hold |

| **Off-target** | **Sequence/ PAM (mismatches)** | **Score** | **Chr** | **Forward primer<br>Reverse primer** | **PCR product** |
|---|---|---|---|---|---|
| 1 | ATGCCAGTCATTCTAGAAGA/ TGG (3) | 2.60 | chr3 | gtggacgacagacatgttgatcc<br>cacaaccaaagccgtccttgaag | 576 |
| 2 | ACCCAATTCTTTCTAGAAGA/ AGG (3) | 1.69 | chr16 | agtgcacactagcactgggatg<br>gaatgtcatgtgagcatgcatggtg | 894 |
| 3 | ACTATAGTTTTTCTAGAAGA/ TGG (3) | 1.58 | chr9 | gggataagcagcagttatctcagg<br>ggcattcagtggtccagataccac | 558 |
| 4 | TCTGCTGTCTTTCTAGAAGA/ TGG (3) | 1.56 | chr14 | ttaggtgggagtgtaggtcagctg<br>ctgaatagccatcagctcctgaacc | 675 |
| 5 | GCTCCAGCCTTTCTAGAACA/ TGG (3) | 1.41 | chr9 | tttcattagggcgcttgctcagg<br>gtgatacagatgtcccactcaagactg | 698 |
| 6 | AGAACATTCTTTCTAGAAGA/ AGG (4) | 0.95 | chr15 | ggatgaatctaccactgctccctctg<br>catgcctcatagcaccatgagatcac | 818 |
| 7 | TGTCCTGGCTTTCTAGAAGA/ TGG (4) | 0.92 | chr5 | gtgctgcacttcagagacaccac<br>acccatgtttcccaggttaggataac | 794 |
| 8 | CCACCAGGCTGTCTAGAAGA/ TGG (4) | 0.91 | chr16 | gcattcatgtggagatgggatggg<br>tctcatagtcgtcagtcacacagcc | 630 |

**Table S3. Sample information.**

| Sample. ID. | Sample name | Sex | Age (days) | Plex/ Mixture | BioReplicate | TMT Channel |
|---|---|---|---|---|---|---|
| 1 | Rbp4Cre APEX+/+ STR group1 (neonate) | M | 5 | 1 | P5_1 | 129C |
| 2 | Rbp4Cre APEX+/+ STR group1 (neonate) | M | 5 | 1 | P5_2 | 129N |
| 3 | Rbp4Cre APEX+/+ STR group1 (neonate) | M | 5 | 2 | P5_3 | 131N |
| 4 | Rbp4Cre APEX+/+ STR group1 (neonate) | F | 5 | 2 | P5_4 | 130N |
| 5 | Rbp4Cre APEX+/+ STR group1 (neonate) | F | 5 | 2 | P5_5 | 132N |
| 6 | Rbp4Cre APEX+/+ STR group2 (early postnatal) | M | 11 | 1 | P11_1 | 130C |
| 7 | Rbp4Cre APEX+/+ STR group2 (early postnatal) | M | 11 | 1 | P11_2 | 133N |
| 8 | Rbp4Cre APEX+/+ STR group2 (early postnatal) | F | 11 | 1 | P11_3 | 132N |
| 9 | Rbp4Cre APEX+/+ STR group2 (early postnatal) | F | 11 | 2 | P11_4 | 131C |
| 10 | Rbp4Cre APEX+/+ STR group2 (early postnatal) | F | 11 | 2 | P11_5 | 133N |
| 11 | Rbp4Cre APEX+/+ STR group3 (preweanling) | M | 18 | 1 | P18_1 | 128C |
| 12 | Rbp4Cre APEX+/+ STR group3 (preweanling) | M | 18 | 1 | P18_2 | 128N |
| 13 | Rbp4Cre APEX+/+ STR group3 (preweanling) | M | 20 | 2 | P18_3 | 127N |
| 14 | Rbp4Cre APEX+/+ STR group3 (preweanling) | F | 18 | 2 | P18_4 | 128C |
| 15 | Rbp4Cre APEX+/+ STR group3 (preweanling) | F | 18 | 2 | P18_5 | 129C |
| 16 | Rbp4Cre APEX+/+ STR group4 (adult) | M | 50 | 1 | P50_1 | 127N |
| 17 | Rbp4Cre APEX+/+ STR group4 (adult) | M | 50 | 1 | P50_2 | 131N |
| 18 | Rbp4Cre APEX+/+ STR group4 (adult) | M | 50 | 1 | P50_3 | 132C |
| 19 | Rbp4Cre APEX+/+ STR group4 (adult) | F | 50 | 2 | P50_4 | 132C |
| 20 | Rbp4Cre APEX+/+ STR group4 (adult) | F | 50 | 2 | P50_5 | 133C |
| 21 | STR CTRL Cre-negative | M | 20 | 1 | STR_negative_1 | 133C |
| 22 | STR CTRL Cre-negative | M | 20 | 1 | STR_negative_2 | 131C |
| 23 | STR CTRL Cre-negative | F | 20 | 2 | STR_negative_3 | 126 |
| 24 | STR CTRL Cre-negative | F | 20 | 2 | STR_negative_4 | 129N |
| 25 | Rbp4Cre H2B CTX | M | 18 | 1 | CTX_H2B_1 | 127C |

| | | | | | | |
|---|---|---|---|---|---|---|
| 26 | Rbp4Cre H2B CTX | M | 18 | 1 | CTX_H2B_2 | 130N |
| 27 | Rbp4Cre H2B CTX | M | 18 | 2 | CTX_H2B_3 | 128N |
| 28 | CTX CTRL Cre-negative | M | 20 | 1 | CTX_negative_1 | 126 |
| 29 | CTX CTRL Cre-negative | M | 20 | 2 | CTX_negative_2 | 127C |
| 30 | CTX CTRL Cre-negative | F | 20 | 2 | CTX_negative_3 | 130C |
| 31 | 5x_STR_REF channel (from P18) | NA | NA | 1 | Norm_1 | 134N |
| 32 | 5x_STR_REF channel (from P18) | NA | NA | 2 | Norm_2 | 134N |

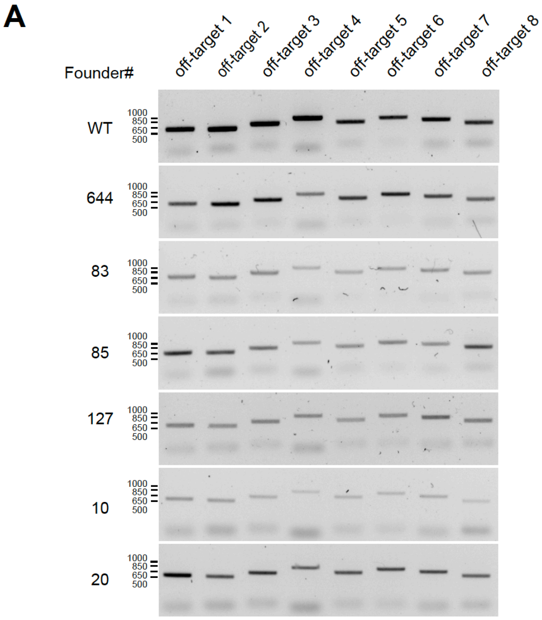


**B**

| Founder# | off-target 1 | off-target 2 | off-target 3 | off-target 4 | off-target 5 | off-target 6 | off-target 7 | off-target 8 |
|---|---|---|---|---|---|---|---|---|
| 644 | 0 | 1 | 0 | 0 | 0 | 0 | 0 | 0 |
| 83 | 0 | 1 | 0 | 0 | 1 | 0 | 0 | 0 |
| 85 | 0 | 1 | 0 | 0 | 0 | 0 | 0 | 1 |
| 127 | 0 | 0 | 0 | 0 | 0 | 0 | 0 | 1 |
| 10 | 0 | 1 | 0 | 1 | 0 | 0 | 0 | 1 |
| 20 | 0 | 1 | 0 | 0 | 0 | 0 | 0 | 0 |

0: 0-1 nucleotide mismatch at the predicted off-target sites
1: double peaks (multiple point mutations) and indels

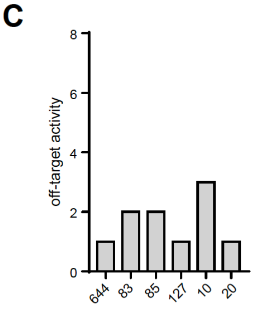


**Figure S1. Cas9 off-target analyses in founder animals.**

**(A)** PCR amplification of genomic regions containing off-target sites predicted by Benchling algorithm across six founder animals including C57BL/6 wild type control.

**(B)** Tabulated off-target activity based on Sanger sequencing of PCR products (0 indicates up to 1 nucleotide mismatch compared to the reference genome, and 1 indicates double peak results or indels).

**(C)** Total number of off-target activity events detected in each founder.

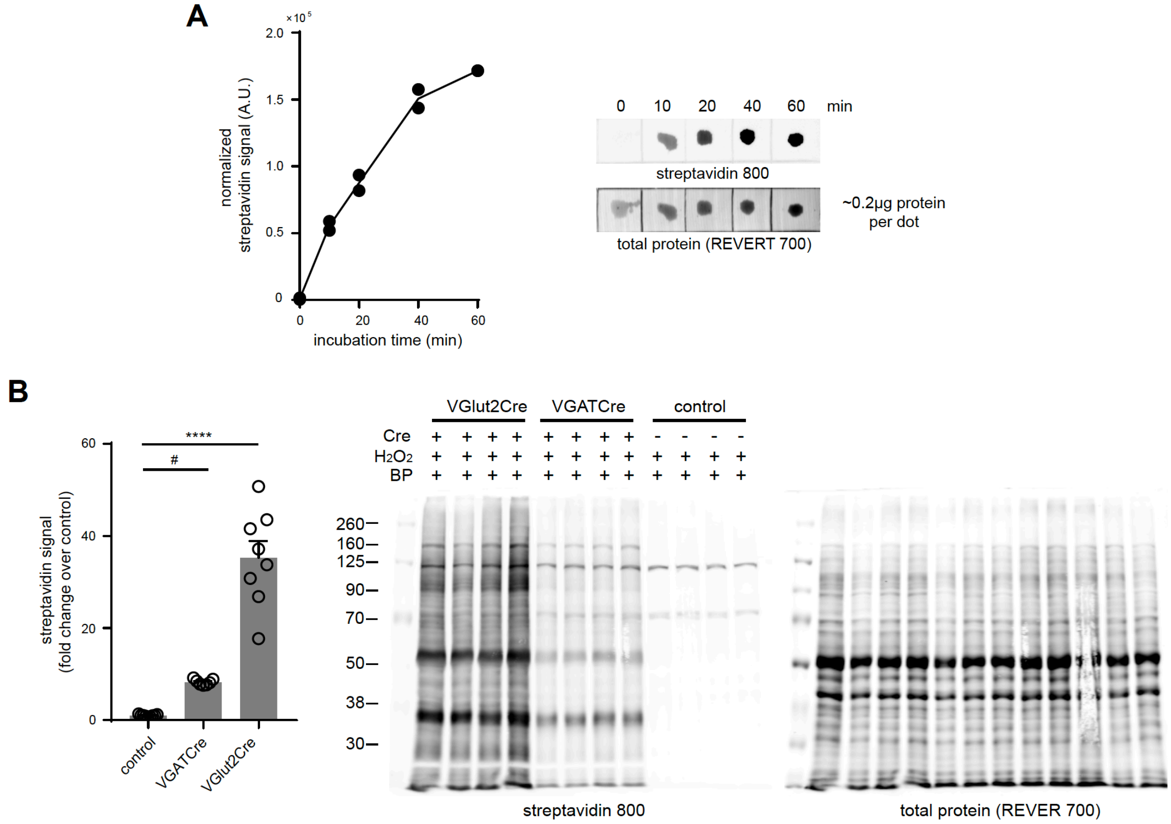


**Figure S2. Optimization of biotin phenol ACSF incubation.**

**(A)** Dot blot analysis of cortical lysates with increasing BP incubation time. *Left*, normalized streptavidin signal plot at time points 0 min, 10 min, 20 min, 40 min and 60 min (n=2 biological replicates). *Right*, representative dot blot.

**(B)** Differential biotinylation in mPFC tissues at 1 hr BP incubation. *Left*, normalized streptavidin signal (one-way ANOVA, $p < 0.0001$, $F(2, 21) = 74.68$, Holm-Sidak's multiple comparison test, VGlut2-Cre vs control, $p < 0.0001$, and VGAT-Cre vs control, $p = 0.0254$) (n= 7 control, 8 VGATCre, 8 VGlut2Cre animals). *Right*, representative streptavidin blot used in quantification.

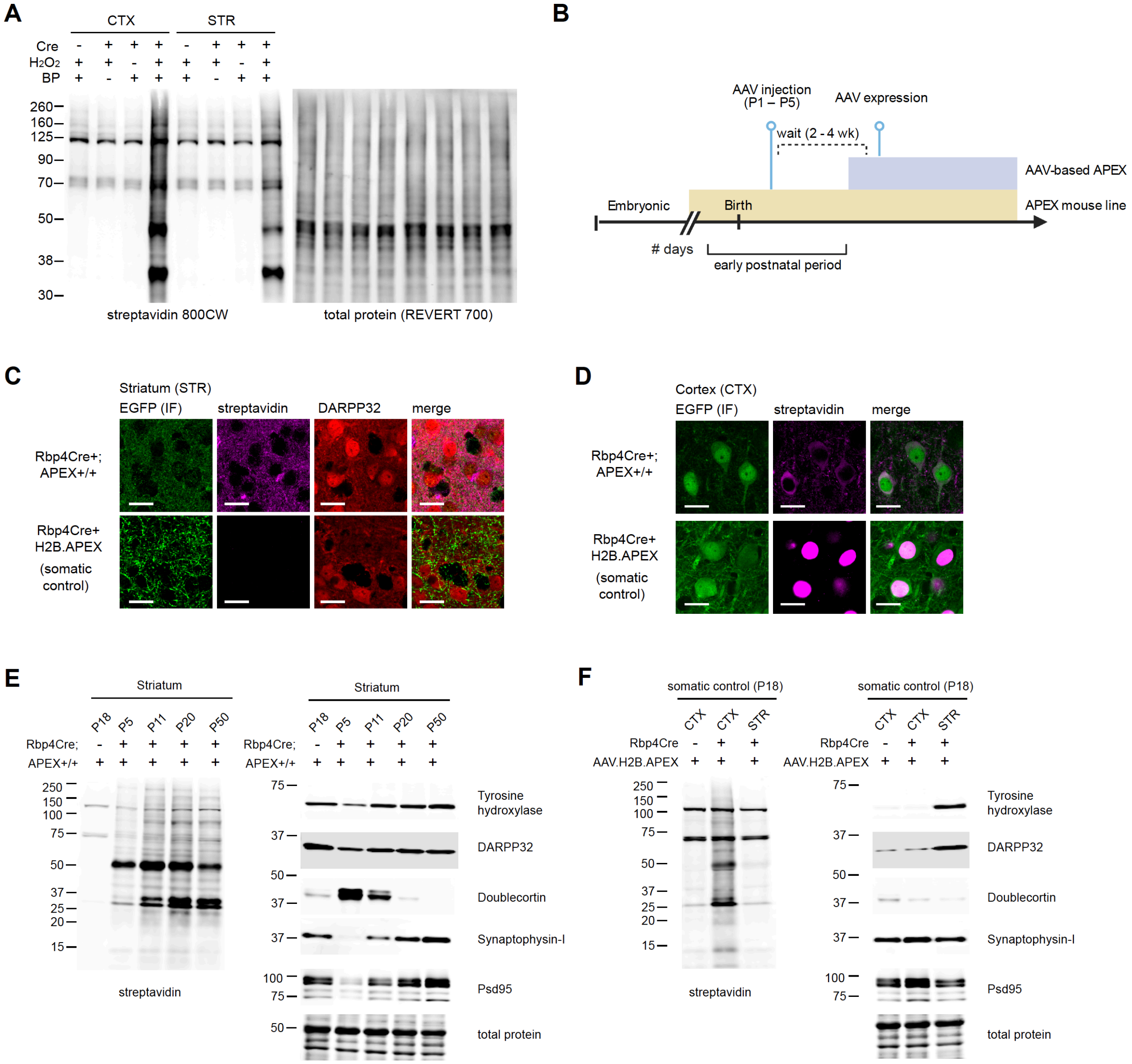


**Figure S3. Cre-dependent biotinylation of Rbp4$^{Cre+}$ corticostriatal projecting neurons.**

**(A)** Western blot analysis of Rbp4$^{Cre+}$;APEX tissue lysates after biotinylation. *Left*, streptavidin blot. *Right*, total protein loading control. Protein labeling by APEX in the cortex (CTX) and striatum (STR) is dependent on Cre, $H_2O_2$, and biotin phenol.

**(B)** APEX expression timeline comparison between AAV and transgenic reporter APEX delivery. Neonatal AAV transduction is usually performed around postnatal days 1-5. Typical wait time before AAV-based expression is approximately 2-4 weeks. Expression from a transgenic reporter usually begins in the embryonic stage, enabling experiments during the early postnatal time window.

**(C)** APEX expression in corticostriatal axons in the dorsal striatum (STR). Rbp4$^{Cre+}$;APEX mouse line expresses APEX.NES in both somata and axons. For APEX-compartment control, Cre-dependent AAV1-H2B.APEX-P2A-EGFP was injected in the cortex of Rbp4$^{Cre+}$ animals to express nucleus-localized H2B.APEX. Confocal images show immunofluorescence for EGFP, streptavidin, and DARPP32 (scale bar: 20 µm). P2A-linked EGFP broadly distributes in axons, while streptavidin labeling confirms proximity labeling of axonal proteins in Rbp4$^{Cre+}$;APEX mouse line but not in H2B.APEX-expressing animals.

**(D)** same as **(C)** for APEX expression in the cortex (CTX). Confocal images show immunofluorescence for EGFP and streptavidin (scale bar: 20 µm). P2A-linked EGFP broadly distributes in somata, while streptavidin labeling confirms proximity labeling of cytosolic and nuclear proteins in Rbp4$^{Cre+}$;APEX mouse line and H2B.APEX-expressing animals, respectively.

**(E)** Western blot analysis of striatal lysates across postnatal development. Acute slices were prepared and biotinylated from Rpb4$^{Cre+}$;APEX animals at the indicated age. Cre-negative APEX reporter line at P18 was used as pull-down (no labeling) control. *Left*, streptavidin blot (same as **Figure 3B**). *Right*, developmental protein markers (Tyrosine hydroxylase, DARPP32, Doublecortin, Synaptophysin-I, and Psd95) and total protein loading control. APEX was expressed in the corticostriatal projections across development confirmed by protein biotinylation.

**(F)** Western blot analysis of striatal and cortical lysates from H2B.APEX-injected Rbp4$^{Cre+}$ animals. Acute slices were prepared and biotinylated from Rpb4$^{Cre+}$;APEX animals at the indicated age. Cre-negative H2B.APEX-injected animal was used as no labeling control. *Left*, streptavidin blot. *Right*, protein markers (Tyrosine hydroxylase, DARPP32, Doublecortin, Synaptophysin-I, and Psd95) and total protein loading control. H2B.APEX primarily labels proteins in the cortex (CTX), with minimal biotinylation in the striatum (STR), serving as somatic protein compartment control.

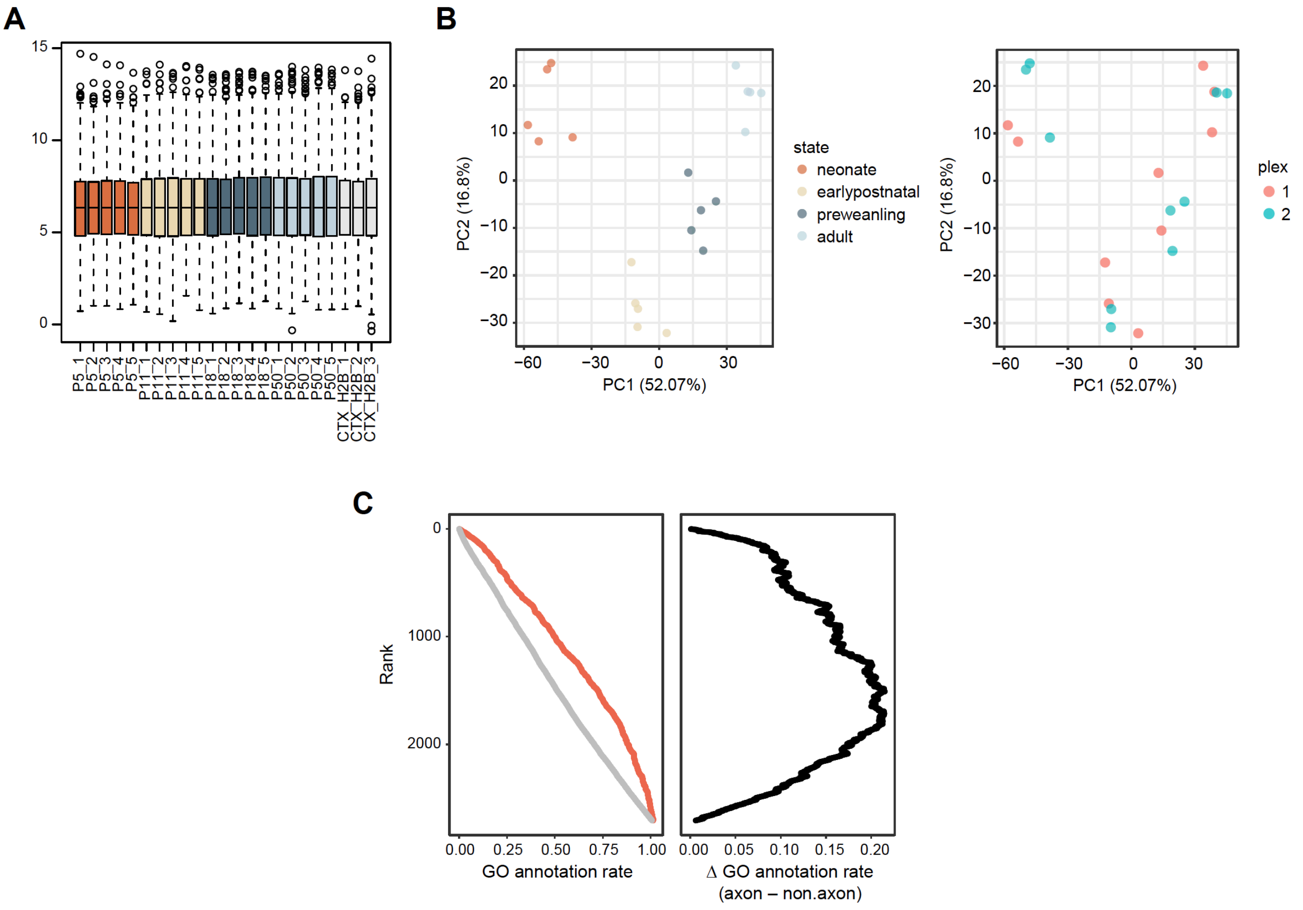


**Figure S4. Quality control plots for the proteomic data.**

**(A)** Boxplot of normalized log2 protein abundance.

**(B)** Principal component analysis of striatal samples.

**(C)** Gene ontology (GO) annotation rate. Proteins were ranked by their log2FC (STR–CTX.H2B). Each protein was marked by GO term, either as axon/presynaptic or not. Cumulative proportion of GO annotation as a function of rank. Red, axon/presynaptic GO terms.

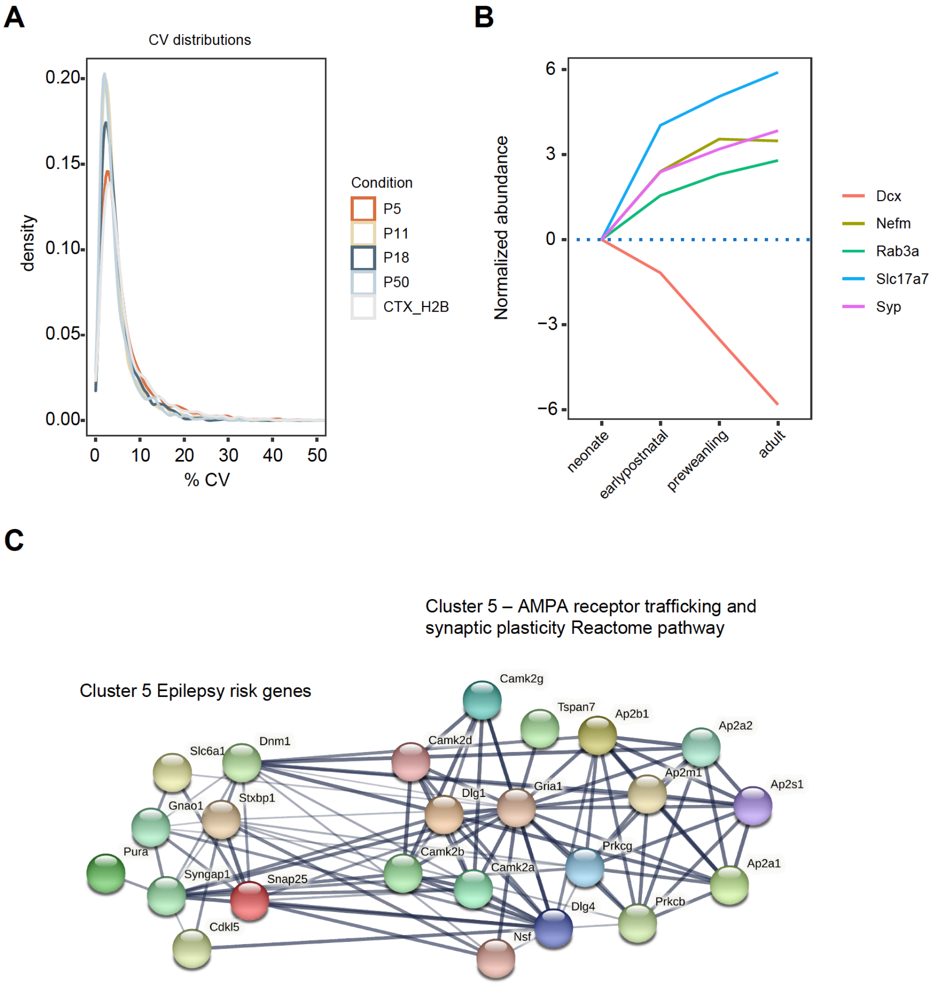


**Figure S5. Sample coefficients of variation (CVs) and expression of selected developmental markers.**

**(A)** Density histogram for sample coefficients of variation (CVs).

**(B)** Developmental trajectories of immature and mature neuronal markers. Doublecortin (*Dcx*), Synaptophysin (*Syp*), vesicular glutamate transporter VGlut1 (*Slc17a7*), intermediate neurofilament (*Nefm*), and vesicle marker *Rab3a*.

**(C)** Proposed relationship between epilepsy (EP) risk genes and cluster 5 biological functions. STRING-DB analysis of EP risk genes in cluster 5 and proteins found in the Reactome pathway for AMPA receptor trafficking and plasticity.

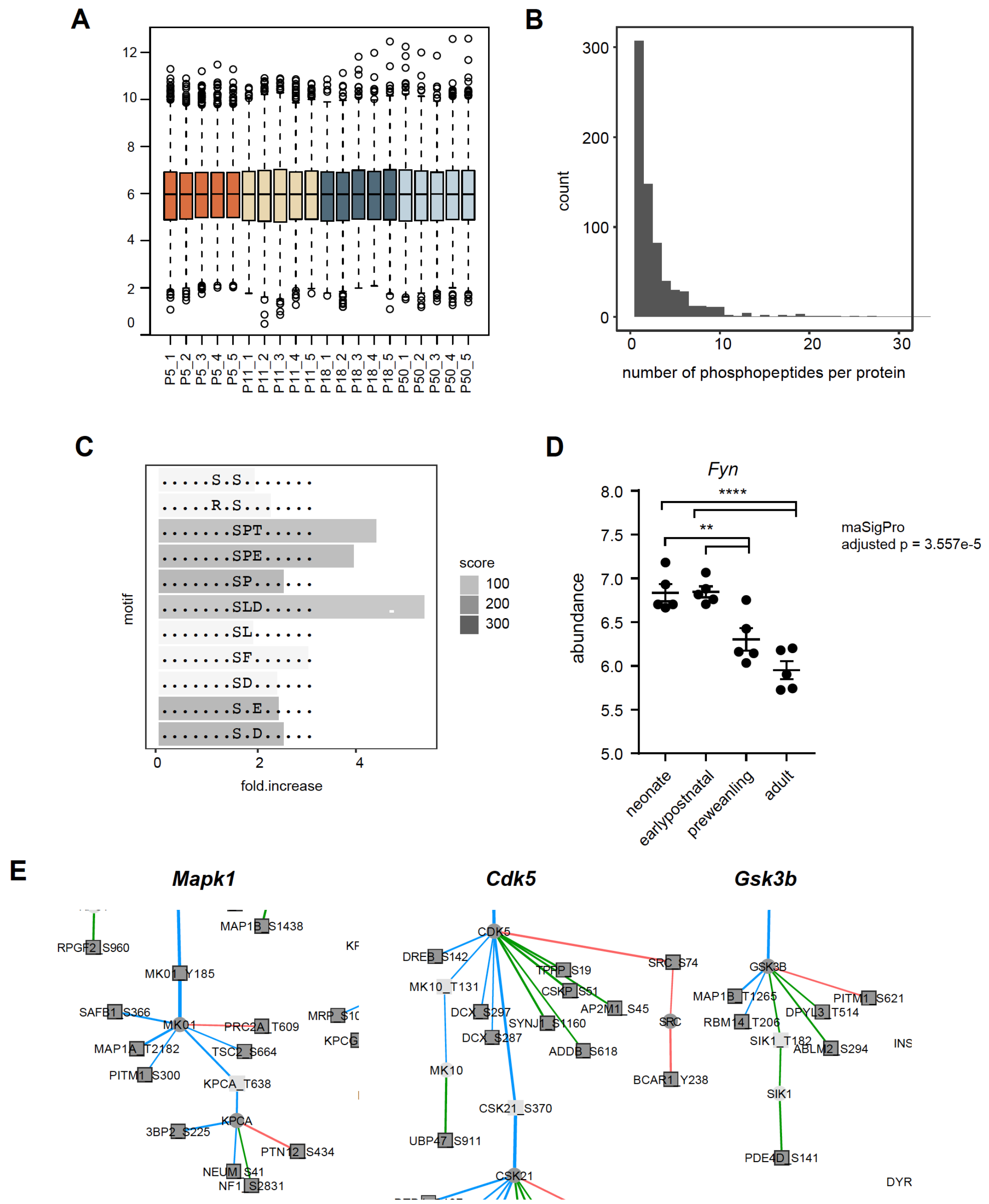


**Figure S6. Phosphopeptide abundance normalization and motif analysis.**

**(A)** Boxplot of normalized log2 phosphopeptide abundance.

**(B)** Number of phosphopeptides per protein. Histogram bin width = 1.

**(C)** Motif-x enrichment analysis.

**(D)** *Fyn* protein log2 abundance. maSigPro adjusted $p = 3.557e-5$. $n = 5$ biological replicates each. One-way ANOVA, $p < 0.0001$, $F(3, 16) = 18.95$, Holm-Tukey's multiple comparison test. Early postnatal vs preweanling, adjusted $p = 0.0078$. Earlypostnatal vs adult, adjusted $p < 0.0001$; neonate vs preweanling, adjusted $p = 0.0088$; neonate vs adult, adjusted $p < 0.0001$.

**(E)** Proline-directed kinases in *Fyn* phosphoregulatory network. Zoom-in for *Mapk1*, *Cdk5*, and *Gsk3b* subnetworks in **Figure 5D**.
